# Bryostatin-1 exploits a developmental signaling vulnerability in MEF2D-rearranged B-cell precursor acute lymphoblastic leukemia

**DOI:** 10.64898/2026.11.12.766137

**Authors:** Quentin Glaziou, Xuekang Qi, Luay Aswad, Isabelle Rose Leo, Frederik Post, Marie Kemkes, Melina Vogt, Owen F.J. Hovey, Georgios Mermelekas, Nona Struyf, Tom Erkers, Katja Pokrovskaja Tamm, Koshi Akahane, Takeshi Inukai, Sanil Bhatia, Janne Lehtiö, Rozbeh Jafari

## Abstract

MEF2D-rearranged (MEF2D-r) B-cell precursor acute lymphoblastic leukemia (BCP-ALL) is a rare molecular subtype associated with high-risk clinical features and frequent relapse and characterized by a distinctive pre-B-cell developmental state and dependence on pre-B-cell receptor (pre-BCR) signaling. Despite these distinctive biological features, MEF2D-r BCP-ALL lacks subtype-directed therapy. Here, we identify a pharmacologically actionable vulnerability to the diacylglycerol (DAG) mimetic bryostatin-1 (Bryo-1). Across 93 BCP-ALL cell lines, MEF2D-r models were enriched for Bryo-1 sensitivity, although responses varied within the subtype and extended to selected non-MEF2D models. Sensitive cells responded at low-nanomolar concentrations with rapid ERK activation followed by loss of pre-BCR-associated components, reduced proliferation, caspase activation and cell death. Pharmacological MEK inhibition markedly attenuated Bryo-1-induced loss of viability, supporting a functional involvement of MEK-ERK signaling. Time-resolved phosphoproteomic and proteomic analyses further revealed coordinated activation of DAG-responsive signaling and subsequent remodeling of the precursor B-cell state. Bryo-1 also altered immunotherapy-relevant surface antigens, including CD19, CD20 and CD22, as part of this broader cell-state transition. Importantly, low-nanomolar activity was reproduced *ex vivo* in MEF2D-r patient-derived xenograft cells. Together, these findings support pharmacological activation of a negative-selection-like developmental checkpoint as a potential therapeutic strategy in responsive MEF2D-r BCP-ALL and provide a framework for biomarker-guided functional patient selection.

## Introduction

B-cell precursor acute lymphoblastic leukemia (BCP-ALL) comprises a heterogeneous group of malignancies defined by distinct initiating genetic alterations and associated biological dependencies^1^. Rearrangements involving the transcription factor MEF2D identify a rare BCP-ALL subtype that is enriched in older children and adolescents and frequently presents with adverse clinical features ^2^. Initial studies associated MEF2D-rearranged (MEF2D-r) BCP-ALL with an increased risk of relapse and inferior survival, despite an often-favorable early response to induction therapy ^3^. More recent analyses have revealed heterogeneity in outcome according to MEF2D fusion partner and clinical context ^4^. Nevertheless, treatment remains based on conventional risk-adapted BCP-ALL regimens rather than therapies directed against the fusion or its downstream biological dependencies, highlighting the need for new therapeutic strategies for this molecular subtype.

MEF2D-r leukemia is characterized by a distinctive developmental state centered on the pre-B-cell receptor (pre-BCR) ^5^. MEF2D fusion proteins promote a transcriptional circuitry that includes genes encoding pre-BCR components and signaling regulators, and pre-BCR expression has been detected in MEF2D-r cell lines and primary leukemia samples. Genetic depletion of pre-BCR components impairs the growth and survival of MEF2D-r cells, indicating that this developmental program contributes directly to leukemia maintenance. During normal B-cell development, antigen-receptor signaling is tightly constrained within a permissive range: insufficient signaling fails to support survival, whereas excessive signaling can cross a pro-apoptotic threshold and engage negative selection ^6^. In developing B cells, this deletional response can involve protein kinase C δ (PKCδ)-and RASGRP-dependent activation of extracellular signal-regulated kinase (ERK), and pharmacological hyperactivation of receptor-proximal signaling has previously been shown to eliminate pre-B-lineage leukemia cells ^7^. These observations suggest that the pre-B-cell identity of MEF2D-r leukemia may create an unusual therapeutic opportunity in which excessive activation, rather than inhibition, of a lineage-defined signaling pathway induces cell death.

Bryostatin-1 (Bryo-1) is a macrocyclic diacylglycerol (DAG) mimetic that binds C1-domain-containing signaling proteins and potently modulates conventional and novel protein kinase C (PKC) isoforms ^8^. Importantly, Bryo-1 has already undergone phase I and phase II clinical evaluation, including in pediatric patients, with a generally manageable safety profile and myalgia reported as the principal dose-limiting toxicity ^9–11^. Our integrative pharmacological and multi-omic screen previously identified Bryo-1 as a candidate therapeutic agent for *MEF2D::HNRNPUL1* BCP-ALL ^12^. In that study, Bryo-1 preferentially reduced the viability of a limited number of MEF2D-r cell lines, activated ERK signaling, and induced cytotoxicity that could be attenuated by MEK or ERK inhibition, suggesting a possible pro-apoptotic PKC-MEK-ERK signaling mechanism. Whether this represented a restricted cell-line observation or a broader vulnerability of MEF2D-r leukemia, however, remained unknown. The pharmacological range of Bryo-1 activity, the molecular sequence linking PKC engagement to cell death, the consequences for pre-B-cell identity, and the extent to which this vulnerability is conserved across MEF2D-r cell lines and patient-derived models have therefore remained unresolved. Bryo-1 has also been reported to alter immunotherapy-relevant B-cell surface antigens, including CD22 ^13–15^, but whether such remodeling occurs as part of the MEF2D-r developmental response has not been established.

Here, we show that Bryo-1 sensitivity is enriched across MEF2D-r BCP-ALL models and occurs at low-nanomolar concentrations. Bryo-1 activates PKC-MEK-ERK associated signaling, followed by attenuation of the pre-BCR-associated state, reduced proliferation and apoptosis, consistent with engagement of a negative-selection-like developmental program. This response extends to MEF2D-r PDX-derived leukemia cells and is accompanied by broader remodeling of the B-cell surface phenotype. Together, these findings support pharmacological activation of a developmental signaling checkpoint as a therapeutic strategy in MEF2D-r BCP-ALL.

## Results

### Bryo-1 reveals a vulnerability enriched in MEF2D-r BCP-ALL

To identify BCP-ALL contexts sensitive to Bryo-1, we examined drug responses across a panel of 93 cell lines spanning diverse genetic backgrounds and fusion subtypes. Responses varied substantially between models, with many lines showing little inhibition and a subset approaching complete inhibition under the screening conditions (**Fig. 1A**). When responses were grouped by molecular subtype, MEF2D-r models generally showed stronger inhibition and greater loss of viability than other subtypes, although sensitivity also occurred outside this group and individual MEF2D-r models remained poorly responsive (**Fig. 1A, B**). This distribution identified MEF2D rearrangement as a context enriched for Bryo-1 sensitivity. We next characterized the duration and cellular consequences of the response. Exposure to 100 nM Bryo-1 for as little as 5 min was followed by a sustained reduction in live-cell numbers during subsequent follow-up (**Fig. 1C**). At 1.5 nM, Bryo-1 reduced ATP-based viability and an independent viability readout in KASUMI-9 and YCUB-4, with increased cytotoxicity and caspase-3/7 activity (**Fig. 1D-G**). KASUMI-9 responded more rapidly than YCUB-4, whereas YCUB-4R showed substantially less loss of viability (**Fig. 1D, E**). Thus, the response in sensitive models included cell death with caspase activation (**Fig. 1F, G**), rather than being explained solely by a reduction in cellular ATP (**Fig. 1D**). Dose-response profiling at days 3 and 6 revealed non-monotonic responses in several models, with stronger inhibition at intermediate concentrations than at the highest concentrations tested (**Fig. S1A**). The position and extent of this response varied between cell lines. These findings established both pronounced sensitivity in a subset of MEF2D-r models and heterogeneity in the concentration dependence and kinetics of the response.

**Figure 1.**
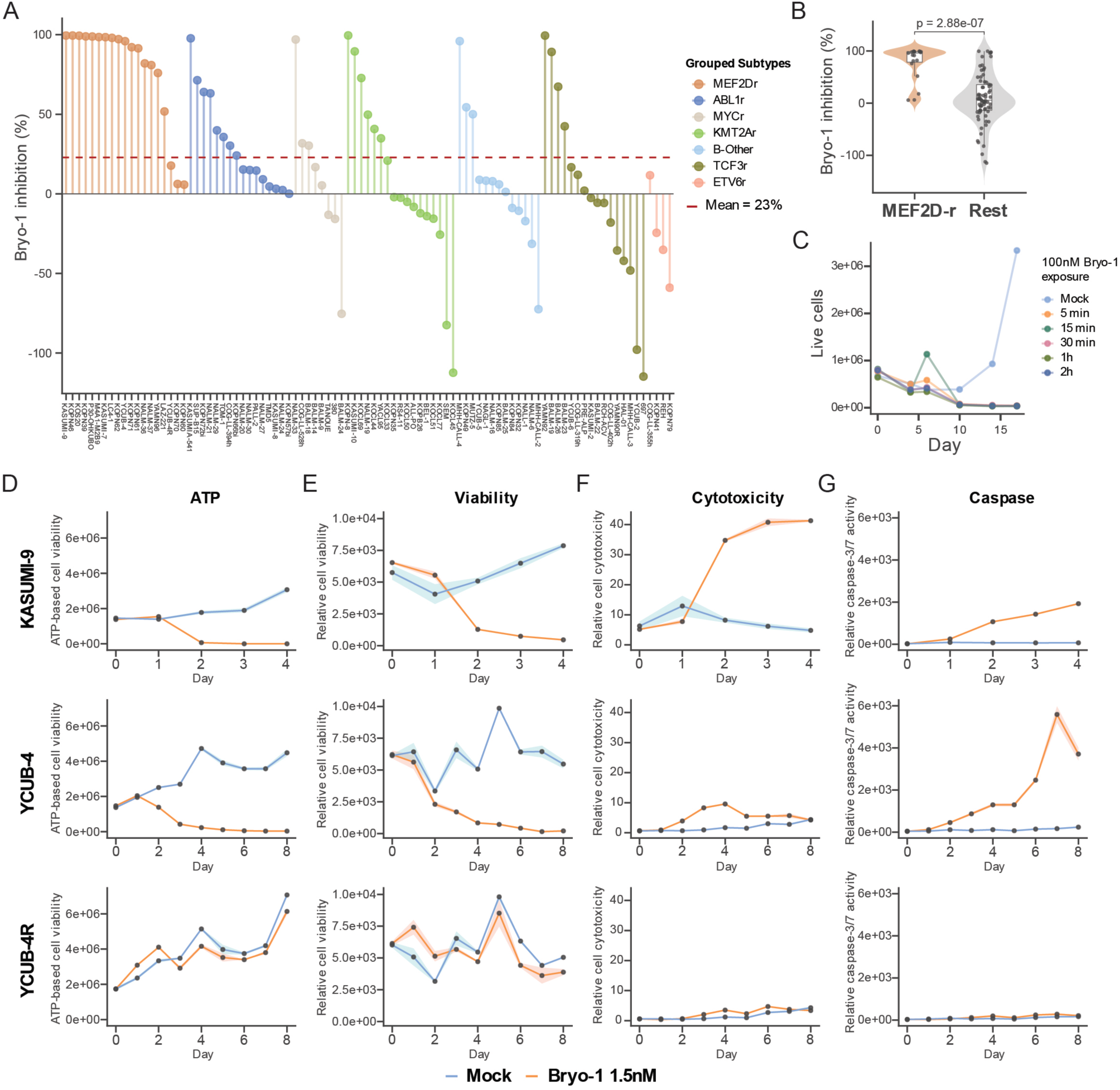
Bryostatin-1 sensitivity is enriched in MEF2D-rearranged BCP-ALL cell lines. **(A)** Inhibition of cell viability across a panel of 93 B-cell precursor acute lymphoblastic leukemia (BCP-ALL) cell lines following 6 days of treatment with 1.5 nM Bryo-1. Cell viability was measured using the CellTiter-Glo assay, and inhibition was calculated relative to the corresponding DMSO-treated controls. Each bar represents one cell line, with colors indicating molecular subtypes. Horizontal lines indicate the mean value of the response distribution. **(B)** Responses from **(A)** comparing MEF2D-r group with the remaining cell lines in the BCP-ALL panel, with individual cell lines overlaid on box plots. Boxes span the interquartile range, central lines indicate the median, and whiskers extend to the most extreme values within 1.5 times the interquartile range. Statistical significance was assessed using two-sided Wilcoxon rank-sum test. **(C)** Live-cell counts following transient exposure to Bryo-1 (n=1). KASUMI-9 cells were treated with 100 nM Bryo-1 for 5, 15, or 30 min, or 1 or 2 h, washed three times with PBS, and resuspended in fresh, drug-free medium. Live cells were counted at the indicated time points using ChemoMetec cell-counting cassettes and compared with mock-treated controls subjected to the same washing procedure. **(D-G)** Time courses of ATP-based viability (**D**), an independent viability readout (**E**), cytotoxicity (**F**), and caspase-3/7 activity (**G**) in KASUMI-9, YCUB-4, and YCUB-4R cells treated with 1.5 nM Bryo-1 or mock control. ATP-based viability was measured using CellTiter-Glo. The additional viability, cytotoxicity, and caspase-3/7 measurements were obtained using the ApoTox-Glo Triplex assay. Data are presented as mean ± SD from three independent experiments; solid lines indicate the mean and shaded bands indicate ± SD.

### Bryo-1 perturbs PKC-associated signaling and induces a MEK-dependent ERK response

To investigate signaling events associated with Bryo-1 sensitivity, we first examined the proteome-wide thermal response to Bryo-1 in KASUMI-9 cells using 2D-TPP ^16,17^. Several PKC isoforms, including PRKCD, PRKCB, PRKCA and PRKCE, together with PRKD2, showed prominent compound-associated thermal shifts (**Fig. 2A**). These proteins contain DAG-responsive C1 domains, consistent with perturbation of this signaling protein family. Additional perturbation of DAG-associated signaling using SAG or the DAG lipase inhibitor RHC 80267 also reduced ATP-based viability in KASUMI-9 cells (**Fig. S1B**). Concentration-and temperature-dependent thermal profiles of PKC proteins, together with CETSA validation of PRKCD, further supported engagement of PKC-associated signaling by Bryo-1 (**Fig. 2B,C**). We next examined the downstream MAPK response. Immunoblotting showed increased ERK1/2 phosphorylation following treatment with 100 nM Bryo-1 in KASUMI-9, YCUB-4 and YCUB-4R cells (**Fig. 2D**). Importantly, ERK activation was also observed at the substantially lower concentration of 1.5 nM, with an increase in the pERK/ERK ratio within 15 min, a peak at approximately 2 h and persistence above baseline at later time points (**Fig. 2E**). Consistent with our previous demonstration that Bryo-1 responses can be attenuated by MEK/ERK inhibition ^12^, trametinib markedly attenuated the Bryo-1-induced reduction in CellTiter-Glo signal in KASUMI-9 and NALM-37 at days 3 and 6 (**Fig. 2F**), showing that this functional dependence is retained under the low-nanomolar Bryo-1 conditions used here. ERK activation was nevertheless also observed in the comparatively poorly responsive YCUB-4R model, indicating that pathway activation alone was not sufficient to determine cytotoxic outcome. We therefore used time-resolved phosphoproteomics to define the broader signaling network engaged downstream of Bryo-1.

**Figure 2.**
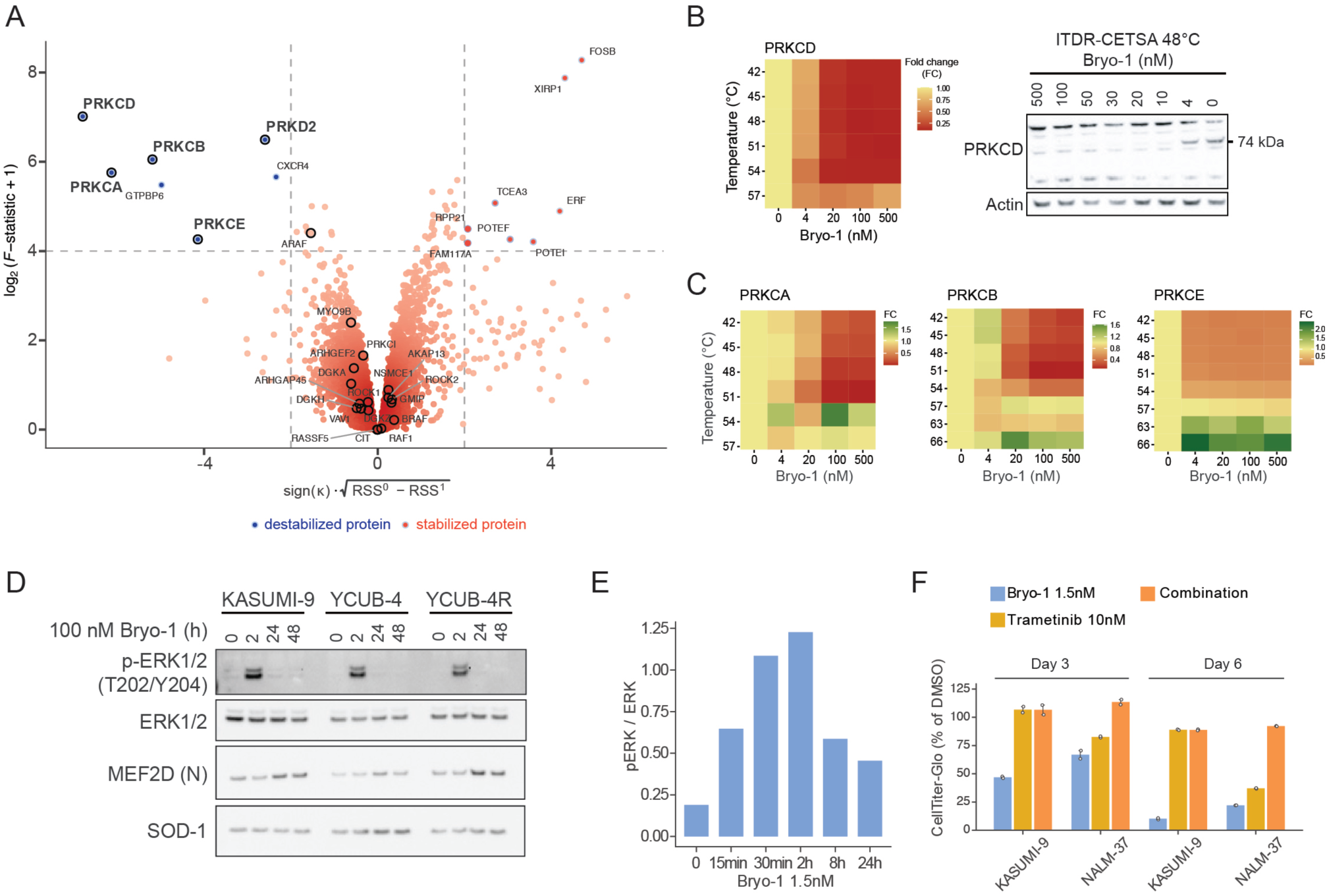
Bryostatin-1 engages PKC-ERK signaling and MEK inhibition counteracts its effect on viability. (**A**) Volcano plot for the 2D-TPP dataset acquired upon treatment of KASUMI-9 cells with Bryostatin-1. Each point represents one protein. The axes show the signed model-comparison score and log2(F-statistic + 1). Proteins containing the Protein kinase C-like, phorbol ester/diacylglycerol-binding domain (PKC_DAG/PE domain, InterPro entry IPR002219) are highlighted with bigger dots. Blue and orange indicate proteins significantly thermally de-stabilized and stabilized by Bryo-1, respectively. (**B**) 2DTPP fold-change (FC) heatmap of soluble PRKCD in KASUMI-9 cells and immunoblot analysis of PRKCD by CETSA at 48°C across a range of Bryo-1 concentrations. The PRKCD band was detected at approximately 74 kDa. Actin served as a loading control. (**C**) 2DTPP FC heatmaps of soluble PRKCA, PRKCB and PRKCE in KASUMI-9 cells. Temperature-and concentration-dependent profiles were measured following 2 h of treatment with Bryo-1 at 0, 4, 20, 100, or 500 nM. Cells were heated at the indicated temperatures from 42 to 66°C, and soluble protein abundance was quantified by mass spectrometry. Missing values at specific temperatures indicate protein abundance below the detection or quantification limit. Values represent fold changes relative to the vehicle-treated sample at the corresponding temperature. Colors indicate relative soluble protein abundance, with yellow and red representing higher and lower values, respectively. (**D**) Immunoblots of phospho-ERK1/2 (Thr202/Tyr204), total ERK1/2, N-terminal MEF2D-reactive proteins, and SOD1 in KASUMI-9, YCUB-4 and YCUB-4R cells treated with 100 nM Bryo-1 for 0, 2, 24, and 48 h. SOD1 served as a loading control (n=1). (**E**) Densitometric quantification of the phospho-ERK1/2-to-total ERK1/2 ratio in KASUMI-9 cells at 0, 15, and 30 min and 2, 8, and 24 h following treatment with 1.5 nM Bryo-1 from western blotting results (n=1). Background-corrected phospho-ERK1/2 and total ERK1/2 signals were each normalized to GAPDH before calculating their ratio. (**F**) ATP-based cell viability in KASUMI-9 and NALM-37 cells at days 3 and 6 following treatment with 1.5 nM Bryo-1 alone, 10 nM trametinib alone or in combination. CellTiter-Glo signals were expressed as percentages of the corresponding DMSO-treated controls. Trametinib was added simultaneously with Bryo-1.

### Bryo-1 rapidly remodels DAG-responsive phosphosignaling and the MEF2D-associated regulatory state

To resolve the signaling events connecting Bryo-1 exposure to the MEK-ERK-dependent phenotype, we performed time-resolved phosphoproteomic profiling of KASUMI-9 cells treated with 100 nM Bryo-1 for 0, 15, 45 and 120 min, with four biological replicates per time point (**Fig. 3A**). The dataset comprised 5,386 class I phosphosites and showed pronounced temporal changes in phosphorylation patterns (**Fig. S2A-C**). A matched total-proteome time course was also generated (**Fig. S2D**). The higher concentration was used to obtain a high-resolution map of the acute signaling response; importantly, ERK activation was independently observed at 1.5 nM Bryo-1, linking the phosphoproteomic perturbation to the pharmacologically active concentration range defined in Figure 2.

**Figure 3.**
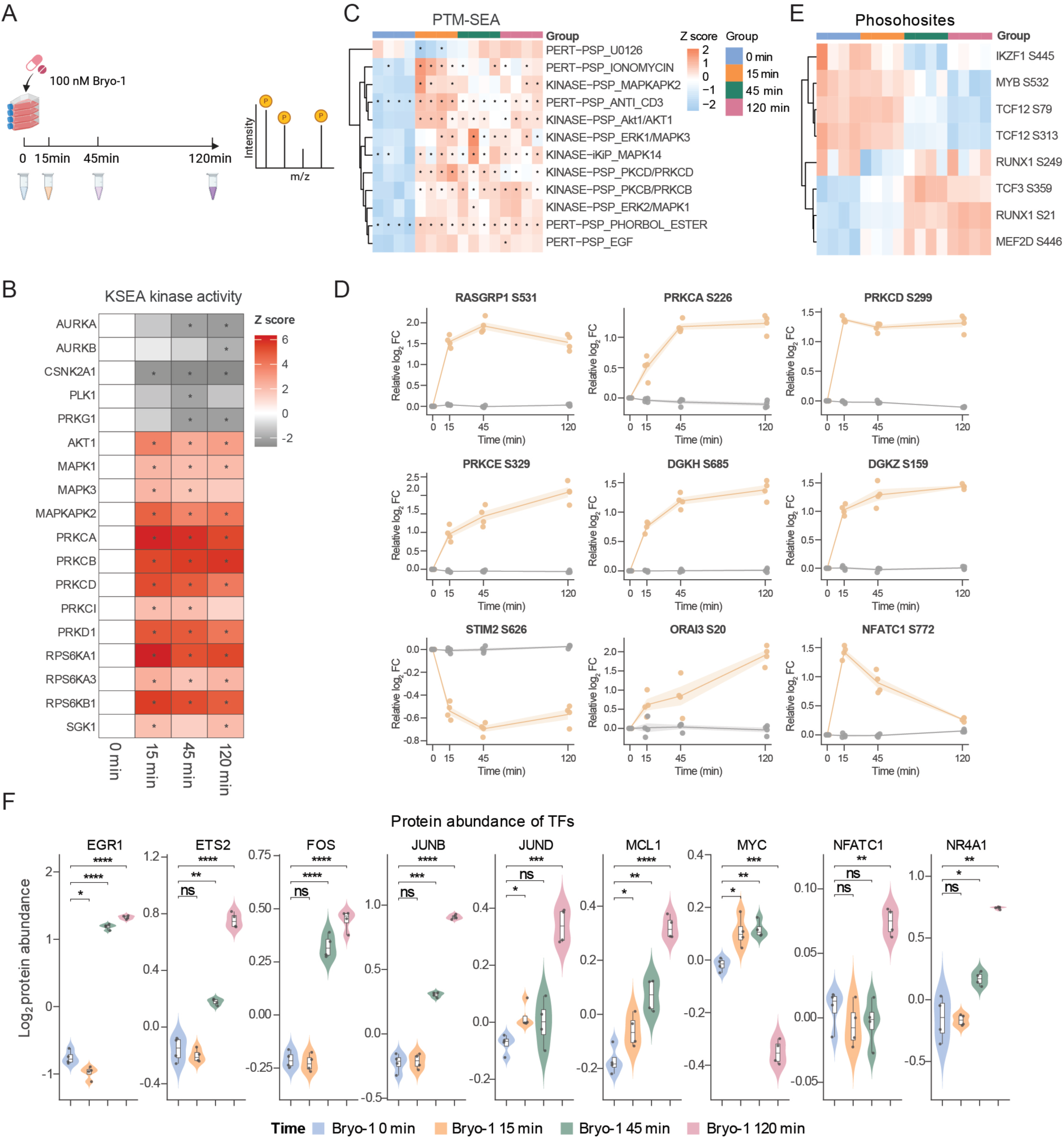
Bryostatin-1 induces rapid PKC/MAPK signaling and transcription-factor phosphoregulation. (**A**) Experimental design and workflow for time-course phosphoproteomic analysis. KASUMI-9 cells were treated with 100 nM Bryo-1 for the indicated time points prior to phosphopeptide enrichment and LC-MS/MS analysis (n=4 biological replicates per time point). (**B**) Kinase activity scores inferred from phosphoproteomic data collected from KASUMI-9 cells at baseline and 15, 45, and 120 min after treatment with 100 nM Bryo-1, using kinase-substrate enrichment analysis with PhosphoSitePlus annotations. Red and gray indicate higher and lower scores, respectively. The *p*-value from KSEA analysis is determined by assessing the one-tailed probability of having a more extreme score than the one measured, followed by a Benjamini-Hochberg FDR correction for multiple hypothesis testing. Kinases with KSEA adjusted p < 0.05 are marked with an asterisk. (**C**) Phosphosite-set enrichment analysis (PTM-SEA) of the time-resolved phosphoproteomic response to Bryo-1. PTM-SEA was performed using phosphosite-specific signatures from PTMsigDB. Rows represent kinase- or perturbation-associated phosphosite signatures and columns represent individual biological replicates grouped by treatment time (0, 15, 45, and 120 min). Enrichment scores were standardized across samples and are displayed as z-scores, with red indicating relative enrichment and blue indicating relative depletion of the corresponding signature. Asterisks indicate statistically significant enrichment/depletion (FDR < 0.02). (**D)** Temporal phosphorylation profiles of selected Bryo-1-responsive phosphosites involved in DAG/PKC signaling, calcium signaling, and downstream signal transduction. Phosphosite abundance is shown as relative log2 fold change over the 0-120 min time course, together with the corresponding total protein abundance where available, illustrating phosphorylation changes occurring independently of changes in protein abundance. Orange indicates phosphosite abundance and gray indicates the corresponding parent protein abundance. Data are presented as mean ± SD from four independent experiments; solid lines indicate the mean and shaded bands indicate ± SD. **(E)** Temporal phosphorylation patterns of selected B-cell and MEF2D-associated transcription factors, including IKZF1, MYB, TCF12, RUNX1, TCF3, and MEF2D, across the 0-120 min Bryo-1 time course. Rows represent individual phosphosites and columns represent replicate samples grouped by treatment time; values are row-scaled phosphosite abundances. **(F)** Protein abundance profiles of EGR1, ETS2, FOS, JUNB, JUND, MCL1, MYC, NFATC1, and NR4A1 in KASUMI-9 cells treated with 100 nM Bryo-1 over the same early time course, expressed as log2 fold changes relative to mean baseline abundance (n=4 biological replicates per time point). Comparisons with baseline were performed using two-sided unpaired Welch’s t-tests: * adjusted P < 0.05, ** < 0.01, *** < 0.001, and **** < 0.0001; ns, not significant.

Kinase-substrate enrichment analysis (KSEA) independently supported increased activity of several PKC-family or DAG-responsive kinases, including PRKCA, PRKCB, PRKCD and PRKD1, together with downstream RPS6KA1, RPS6KB1 and MAPKAPK2 activity (**Fig. 3B**). Phosphosite-set enrichment analysis (PTM-SEA) revealed a rapid and coordinated response involving DAG/C1-responsive and MAPK-associated signaling programs (**Fig. 3C, Fig. S2E**). Among the prominent signatures were those associated with PRKCA and PRKCD, ERK1/2, downstream RSK and MAPKAPK2 signaling, together with a strong phorbol-ester-associated phosphoproteomic signature. Ca²⁺-linked kinase signatures were also altered. The same overall signaling structure was retained when phosphopeptide measurements were corrected for matched parent-protein abundance, indicating that the acute response predominantly reflected changes in phosphorylation state rather than corresponding changes in protein abundance.

At the level of individual phosphoproteins, Bryo-1 induced rapid, time-dependent phosphorylation changes across several components positioned within or adjacent to DAG-responsive signaling (**Fig. 3D**). These included RASGRP1 S531 and sites in the conventional and novel PKC isoforms PRKCA S226, PRKCD S299 and PRKCE S329. Phosphoregulation also extended to proteins controlling DAG/phosphatidic-acid homeostasis, including DGKH S685 and DGKZ S159, and to Ca²⁺-linked signaling components including STIM2 S626, ORAI3 S20 and NFATC1 S772. NFATC1 showed additional site-specific regulation, with phosphorylation at S237 and S887 decreasing over the acute time course without a corresponding decrease in total NFATC1 abundance (**Fig. S2F**). NFATC1 S237 lies within the N-terminal phosphoregulatory region of NFATC1, where multisite phosphorylation contributes to control of NFAT localization and activity ^18–20^. Together, these trajectories indicate that the Bryo-1 response extends beyond an individual PKC isoform and engages a broader network linking C1-domain signaling, lipid second-messenger regulation and Ca²⁺-responsive transcriptional control. RASGRP1 S531 was particularly notable because RASGRP1 provides a direct C1-domain link between DAG-responsive signaling and RAS activation; however, S531 is not currently established as a PKC substrate or functionally annotated phosphosite, and its regulation therefore identifies RASGRP1 as part of the acute phosphoregulatory response rather than demonstrating a specific kinase-substrate relationship.

Because MEF2D-r leukemia is maintained by a lineage-restricted transcriptional program ^5^, we next asked whether the acute signaling response extended to regulators of the MEF2D/pre-B-cell state. Bryo-1 induced coordinated phosphorylation changes in multiple transcriptional regulators, including MYB S532, RUNX1 S21 and S249, TCF3 S359, TCF12 S79 and T313, IKZF1 S445 and MEF2D S446 (**Fig. 3E**). Independent phosphopeptide features mapping to MEF2D S446 showed concordant regulation. The MEF2D S446 phosphopeptide maps to canonical MEF2D and does not distinguish wild-type from fusion-derived MEF2D. Thus, the early response was not restricted to canonical cytoplasmic signaling proteins but reached transcription factors central to B-cell identity and the regulatory context in which the MEF2D fusion operates.

This phosphoregulatory response was followed by a rapid immediate-early protein response. EGR1, ETS2, FOS, JUNB and NR4A1 levels increased within 45-120 min, with particularly strong induction of EGR1 levels, whereas MYC levels declined by 120 min after a modest early increase (**Fig. 3F**). Global clustering of the matched early proteome likewise revealed time-associated abundance changes (**Fig. S2G**). These changes place activation of immediate-early transcriptional programs downstream of the acute phosphosignaling response and precede the more extensive loss of the precursor B-cell and proliferative state observed at later time points.

Together, these data position Bryo-1 as a modulator of a broader DAG-responsive signaling state rather than of a single PKC isoform. Acute PKC- and MAPK-associated phosphosignaling was accompanied by phosphoregulation of RASGRP1, DAG/PA- and Ca²⁺-linked proteins and multiple MEF2D/B-cell regulatory factors, followed by induction of immediate-early transcriptional regulators. The preservation of these responses after accounting for matched parent-protein abundance further supports a genuine signaling response. We next asked how this acute molecular program was translated into the later proteomic and phenotypic transition of the leukemia cells.

### Bryo-1 induces a phenotype consistent with a negative-selection-like response

We next asked how the early signaling response related to later changes in the pre-BCR-associated phenotype and proliferation. CD179a, corresponding to the surrogate light-chain component VPREB1, decreased within the first day of treatment across the eight models examined (**Fig. 4A-C**). Low protein levels persisted in most lines, whereas CD179a levels in YCUB-4R recovered over subsequent days (**Fig. 4A, B**). Total SYK abundance also decreased by 24 h in KASUMI-9, YCUB-4 and YCUB-4R (**Fig. 4D**). The phosphorylated ZAP70 (pZAP70)/ZAP70 ratio showed a more complex time course, with an initial reduction followed by recovery, particularly in YCUB-4 and YCUB-4R (**Fig. 4E**). Proteomic analysis revealed accompanying changes in the machinery supporting the precursor B-cell state. Global analysis of the 6- and 24-h proteome confirmed treatment-associated changes in protein abundance and coordinated temporal remodeling (**Fig. S3A-D**). At 24 h, the response included lower abundance of IGLL1, VPREB1, BLNK and IL7R, together with reduced MYC and BCL2 (**Fig. 4F, G**). PTPN6 and CBL increased, as did EGR2, CDKN1A and the apoptosis-associated proteins CASP3, CASP8 and CASP9. EGR1 showed an earlier increase at 6h. The late proteome also revealed broader surface remodeling, with increased CD19, CD22, MS4A1/CD20 and ICAM1 and reduced MME/CD10, particularly at 24 h (**Fig. 4G**). These changes connected loss of pre-BCR components with increased abundance of negative regulators of signaling and proteins associated with growth restraint and apoptosis. Consistent with this response, Ki-67 fluorescence decreased over three days in most models, while YCUB-4R showed a partial recovery at the final time point (**Fig. 4H**). Proteome enrichment analyses also identified reductions in cell-cycle, G1/S-transition and DNA-repair-associated categories (**Fig. S3E,F**). Taken together with the early ERK response, its pharmacological antagonism by MEK inhibition and the caspase-associated loss of viability, these findings support a negative-selection-like phenotype characterized by attenuation of the precursor signaling state, broader cell-surface remodeling, and reduced proliferative activity.

**Figure 4.**
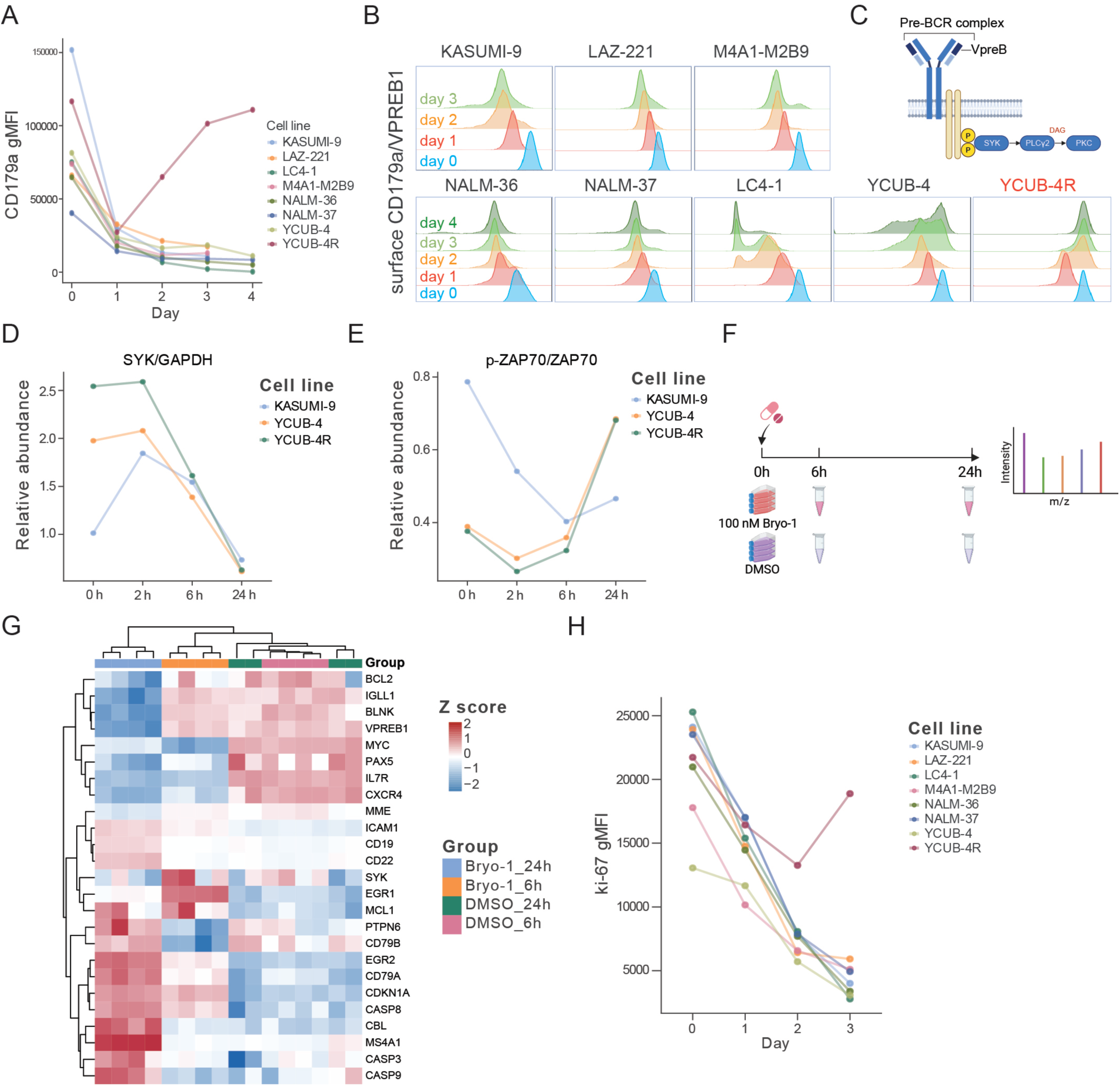
Bryostatin-1 reduces pre-BCR components and proliferation-associated markers. (**A-B**) Time course of CD179a/VPREB1 levels in KASUMI-9, LAZ-221, LC4-1, M4A1-M2B9, NALM-36, NALM-37, YCUB-4, and YCUB-4R cells following treatment with 1.5 nM Bryo-1 (n=1). Protein levels were measured by flow cytometry and is presented as gMFI over days 0-4, with day 0 corresponding to the pretreatment baseline (**A**). Corresponding fluorescence distributions are shown for each cell line (**B**). Histogram colors indicate sampling time: cyan, day 0; red, day 1; orange, day 2; light green, day 3; and dark green, day 4. (**C**) Schematic model of the pre-BCR complex and downstream pathway. Densitometric quantification of total SYK abundance (**D**) and the phospho-ZAP70-to-total ZAP70 ratio (**E**) in KASUMI-9, YCUB-4, and YCUB-4R cells at 0, 2, 6, and 24 h after treatment with 1.5 nM Bryo-1. Total SYK signals were normalized to GAPDH. Phospho-ZAP70 and total ZAP70 signals were each normalized to GAPDH before calculating their ratio. Red, yellow-green, and purple indicate KASUMI-9, YCUB-4, and YCUB-4R, respectively (n=1). (**F**) Experimental design and workflow for quantitative proteomic analysis. KASUMI-9 cells were treated with DMSO or 100 nM Bryo-1 for 6 h or 24 h prior to LC-MS/MS analysis. (**G**) Heat map of selected proteins associated with pre-BCR signaling, precursor B-cell identity, growth regulation, apoptosis, and cell-surface remodeling in KASUMI-9 cells treated with 100 nM Bryo-1 or DMSO for 6 or 24 h. Rows represent proteins and columns represent individual samples, with four samples per condition. Log2 protein intensities were standardized within each row to generate z-scores. Red and blue indicate relatively higher and lower protein abundance, respectively. Dendrograms show hierarchical clustering of proteins and samples, and the upper annotation identifies treatment and sampling time. (**H**) Time course of intracellular Ki-67 levels in the cell lines listed in (A) following treatment with 1.5 nM Bryo-1. Ki-67 was measured by flow cytometry after fixation and permeabilization and is presented as gMFI over days 0-3 (n=1).

### Bryo-1 remodels surface antigens and maturation-associated markers

Because the late proteome indicated broader remodeling of the leukemia-cell surface, we next examined the corresponding cell-surface phenotype by flow cytometry. By day 2, CD19 fluorescence increased to varying degrees across the eight models examined (**Fig. 5A**), while CD22 increased in most lines and remained low in LC4-1 (**Fig. 5B**). CD20 induction was particularly marked in several models, with increases in both CD20 fluorescence and the fraction of CD20-positive cells (**Fig. 5C, D**). Representative histograms confirmed these changes relative to time matched DMSO controls (**Fig. 5E**). The response remained heterogeneous, with comparatively limited CD20 induction in KASUMI-9 and minimal induction in LC4-1 (**Fig. 5C-E**). Antigen remodeling was accompanied by changes in additional surface markers. CXCR4/CD184 fluorescence and the fraction of CD184-positive cells fell sharply within one day (**Fig. 5F, G**), while CD10 and CD24 decreased in several models (**Fig. 5H-I**). Consistent with these flow-cytometric changes, proteomics data showed increased MS4A1/CD20 and decreased CXCR4, IL7R and CD38 at 24 h (**Fig. S3C,D**). CD43 and CD99 followed heterogeneous trajectories, indicating that the response did not impose a uniform change across all phenotypic markers (**Fig. S3G**). The loss of precursor-associated features together with CD20 acquisition was compatible with a maturation-associated shift in cell state. Analysis at 6 and 24 h showed changes in BATF, IRF7 and STAT1 abundance in KASUMI-9 (**Fig. S3B,C**), consistent with regulatory remodeling. Notably, YCUB-4R retained substantial induction of CD19, CD20 and CD22 despite its limited loss of viability and recovery of CD179a and Ki-67. Surface-antigen remodeling could therefore occur in a model that was poorly responsive to the cytotoxic effect of Bryo-1, separating these two features of the response (**Figs. 1D**, **4A, H** and **5A-E**).

**Figure 5.**
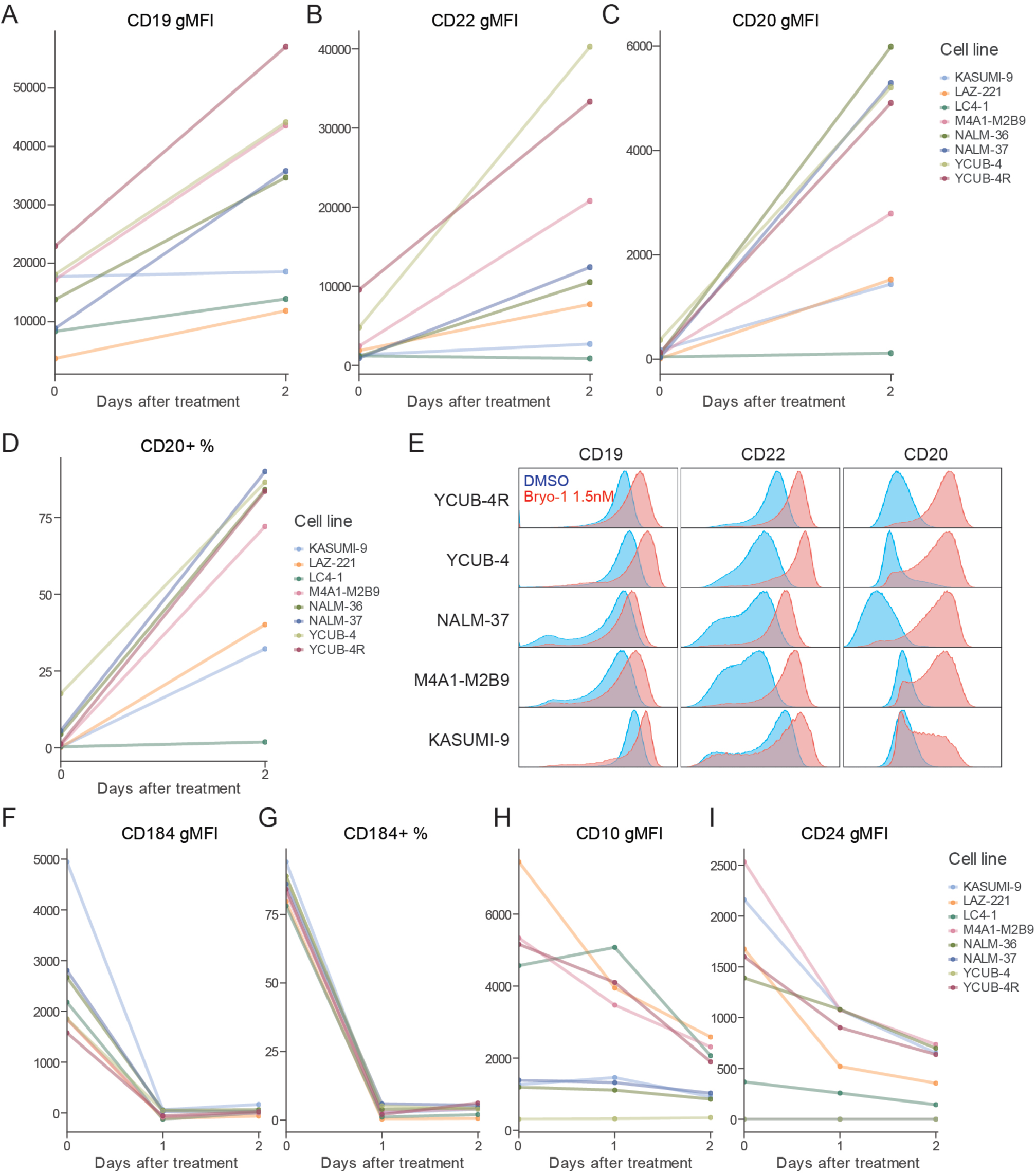
Bryostatin-1 remodels surface antigens and maturation-associated markers. (**A-D**) Surface levels of CD19, CD22, and CD20 in KASUMI-9, LAZ-221, LC4-1, M4A1-M2B9, NALM-36, NALM-37, YCUB-4, and YCUB-4R cells at baseline and 2 days after treatment with 1.5 nM Bryo-1. Protein levels were assessed by flow cytometry and is presented as gMFI for CD19 (**A**), CD22 (**B**), and CD20 (**C**), together with the percentage of CD20-positive cells (**D**). Colors identify individual cell lines, and lines connect baseline and day-2 measurements within each cell line (n=1). (**E**) Representative flow-cytometry distributions of surface CD19, CD22, and CD20 in KASUMI-9, M4A1-M2B9, NALM-37, YCUB-4, and YCUB-4R cells after 2 days of treatment with 1.5 nM Bryo-1 or DMSO. Cyan and red distributions represent DMSO-treated control and Bryo-1-treated cells, respectively. (n=1) (**F-I**) Time courses of surface CXCR4/CD184 (**F**) or the percentage (**G**) of positive cells, CD10 (**H**), and CD24 levels (**I**) over days 0-2 following treatment with 1.5 nM Bryo-1 in the cell lines listed in (**C**). (n=1)

### Bryostatin-1 induces heterogeneous cytotoxic and immunophenotypic responses in MEF2D-rearranged PDX models

To determine whether the Bryo-1 response observed in cell lines extended to patient-derived leukemia models, we examined human leukemia cell enriched preparations from three independent *MEF2D::BCL9* PDXs. Bryo-1 produced marked concentration- and time-dependent reductions in viable-cell signal in SJBALL020667 and SJBALL020987, with substantially greater effects at day 6 than at day 3 (**Fig. 6A,B**) consistent with the observations in cell line models. In contrast, SJBALL021005 showed little reduction in viability across the concentration range tested and was therefore comparatively resistant to the cytotoxic effects of Bryo-1 (**Fig. 6C**). These findings demonstrate substantial inter-model heterogeneity in the magnitude of the viability response despite the shared MEF2D-r genotype. To further assess the contribution of MEK-ERK signaling in patient-derived models, we examined Bryo-1 responses in bulk, unsorted SJBALL020667 PDX derived cells. Co-treatment with 0.5 µM MEK inhibitor U0126 attenuated the Bryo-1-induced reduction in ATP-based viability signal at day 3 (**Fig. S4**), consistent with the MEK inhibitor antagonism observed in established MEF2D-r cell lines (**Fig. 2F**). We next examined whether Bryo-1 nevertheless induced phenotypic remodeling in these models. Expression of the pre-B-cell-associated marker CD179a decreased after treatment in the responsive PDXs, although the magnitude and kinetics differed between models (**Fig. 6D**). CXCR4 was rapidly reduced in all three PDXs, followed by variable recovery at later time points (**Fig. 6E**). Ki-67 also changed in a model-dependent manner, with the clearest reduction observed in SJBALL020987, whereas SJBALL020667 remained comparatively stable and the resistant SJBALL021005 showed a different temporal profile (**Fig. 6F**). CD43 and CD10 similarly displayed heterogeneous trajectories, indicating that Bryo-1 does not impose an identical differentiation phenotype across individual PDXs (**Fig. 6G,H**). Bryo-1 also altered immunotherapy-relevant B-lineage surface antigens, but these responses were not uniformly coupled to cytotoxic sensitivity. CD19 increased in all three PDXs, with particularly strong induction in SJBALL021005 despite its limited viability response (**Fig. 6I**). CD20 induction was more heterogeneous: expression increased clearly in SJBALL020667 and SJBALL021005 but remained low in SJBALL020987 despite its sensitivity to Bryo-1 (**Fig. 6J**). CD22 likewise increased in multiple models, again with the strongest induction observed in SJBALL021005 (**Fig. 6K**). Thus, the magnitude and spectrum of antigen remodeling differed between PDXs and did not simply mirror the extent of Bryo-1-induced loss of viable-cell signal. Together, these data extend the Bryo-1-induced state transition to patient-derived MEF2D-r BCP-ALL while revealing inter-model heterogeneity in its downstream phenotypic consequences. In particular, the preservation of strong surface-antigen remodeling in a comparatively cytotoxic-response-resistant PDX, together with incomplete CD20 induction in one sensitive model, indicates that cytotoxicity, differentiation-associated remodeling and modulation of therapeutic surface antigens can represent partially separable outputs of Bryo-1 exposure.

**Figure 6.**
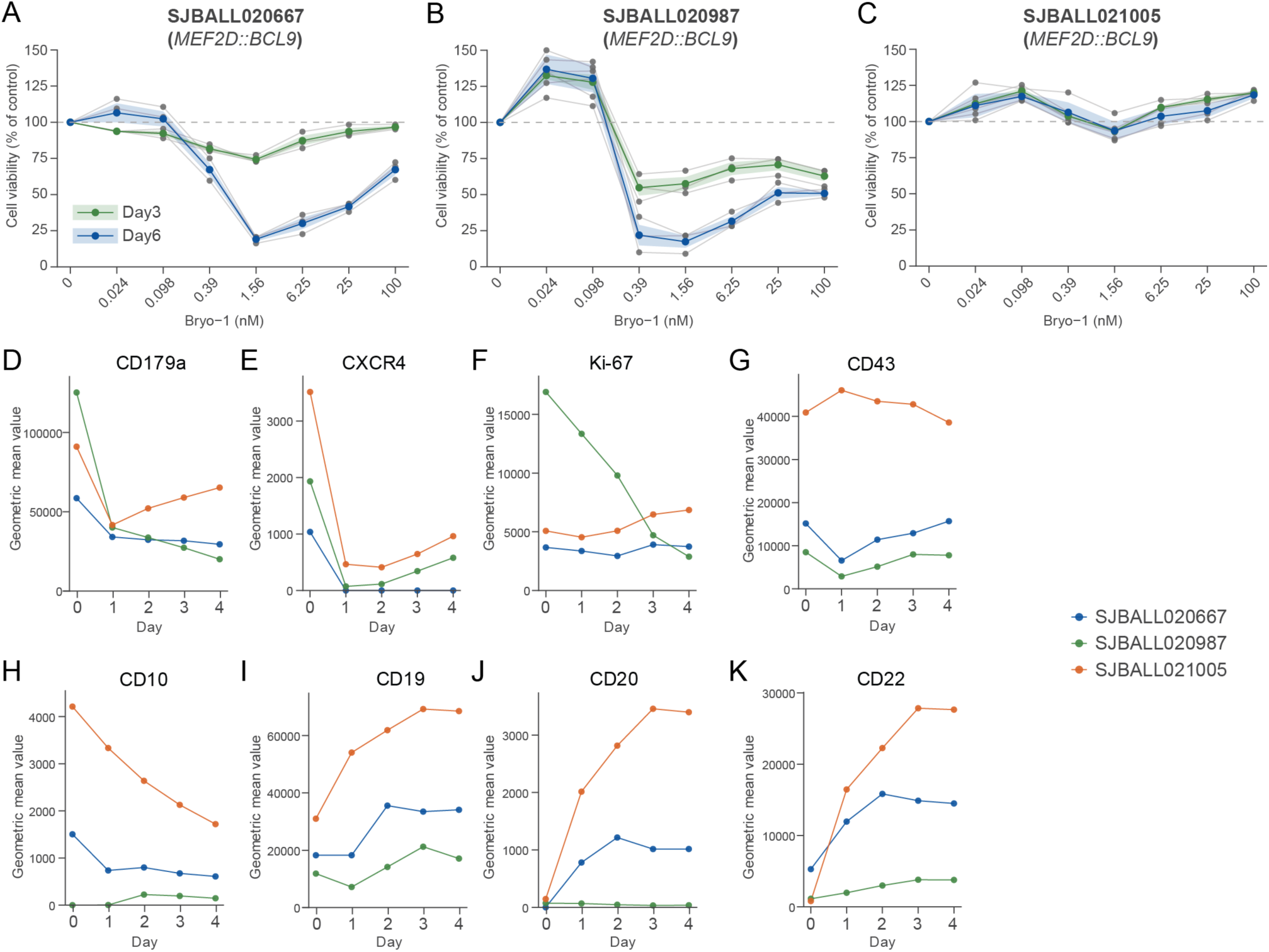
Bryostatin-1 induces heterogeneous cytotoxic and immunophenotypic remodeling in MEF2D-rearranged PDX models. **(A-C)** Concentration-response profiles of three independent *MEF2D::BCL9* patient-derived xenograft (PDX) models, SJBALL020667, SJBALL020987 and SJBALL021005, following Bryo-1 treatment. Cell viability was measured on day 3 (green) and day 6 (blue) and is shown relative to vehicle-treated controls. SJBALL020667 and SJBALL020987 displayed marked concentration- and time-dependent reductions in viable-cell signal, with maximal effects within the low-nanomolar concentration range, whereas SJBALL021005 showed comparatively limited cytotoxic response across the tested concentration range. Gray points indicate individual technical wells (n=3); colored lines and shaded bands indicate mean ± SD. **(D-H)** Time-resolved flow-cytometric analysis of differentiation-, precursor-state- and proliferation-associated markers following 1.5 nM Bryo-1 exposure were obtained once for each PDX model. Geometric mean fluorescence intensity is shown for **(D)** CD179a, **(E)** CXCR4, **(F)** Ki-67, **(G)** CD43 and **(H)** CD10 from day 0 to day 4. Bryo-1 induced changes in multiple phenotypic markers across the three PDX models, including reduction of the pre-B-cell-associated marker CD179a and rapid modulation of CXCR4, although the magnitude and persistence of individual responses differed between models. **(I-K)** Time-dependent modulation of immunotherapy-relevant B-lineage surface antigens following 1.5 nM Bryo-1 exposure. Geometric mean fluorescence intensity is shown for **(I)** CD19, **(J)** CD20 and **(K)** CD22. CD19 and CD22 increased in multiple PDX models, whereas CD20 induction was more heterogeneous. Notably, substantial antigen remodeling was also observed in the comparatively cytotoxic-response-resistant SJBALL021005 model, indicating that surface-antigen modulation and loss of viable-cell signal can be partially uncoupled. Colors in D-K indicate SJBALL020667 (blue), SJBALL020987 (green) and SJBALL021005 (orange).

## Discussion

This study identifies a vulnerability to Bryo-1 that is enriched in MEF2D-r BCP-ALL and connects its cytotoxic activity to broader remodeling of the leukemic cell state. Across 93 cell lines, MEF2D rearrangement marked a particularly responsive group, although sensitivity extended beyond this subtype and varied within it. In sensitive models, early ERK activation was followed by attenuation of the pre-BCR-associated program, reduced proliferation and caspase-associated cell death. Bryo-1 also increased the surface levels of antigens used for immunotherapy, including in a model with limited cytotoxic sensitivity. Together, these findings support the development of Bryo-1 as a direct antileukemic strategy in selected BCP-ALL contexts, while identifying surface-antigen remodeling as a potentially complementary translational consequence of the drug-induced cell-state transition.

The association between ERK activation and leukemia cell death places these findings within the biology of developmental selection. Developing B cells must maintain antigen-receptor signaling within a range compatible with survival, and excessive signaling can engage deletional responses. Limnander and colleagues linked PKCδ and RASGRP to a proapoptotic ERK threshold during B-cell development ^6^, while studies in B-cell acute lymphoblastic leukemia (B-ALL) showed that enforced proximal signaling or disruption of ERK negative feedback can expose a retained susceptibility to negative selection ^7,21^. Our results extend this therapeutic principle to a pharmacologically accessible response enriched in MEF2D-r leukemia. Rapid ERK activation, antagonism by MEK inhibition, induction of the antigen-receptor-responsive transcription factor NR4A1, and subsequent apoptosis support a negative-selection-like mechanism. This response was accompanied by loss of the pre-BCR surrogate light-chain components VPREB1/CD179a and IGLL1, consistent with broader remodeling of the pre-B-cell developmental program. This interpretation concerns the persistence of a signaling vulnerability associated with developmental checkpoints; the experiments do not establish antigen-dependent autoreactivity or complete reconstruction of physiological negative selection.

MEF2D-r leukemia provides a biologically coherent setting for this response. MEF2D fusion proteins participate in a transcriptional circuitry that sustains pre-BCR signaling and the precursor B-cell state, and inhibition of this circuitry impairs leukemia maintenance ^5^. Our findings suggest that excessive activation of downstream signaling offers another way to destabilize this state. The temporal sequence is informative: induction of EGR1, AP-1 family proteins and NR4A1 precedes reductions in pre-BCR components, MYC and BCL2, together with increased CDKN1A and apoptosis-associated proteins. Increased PTPN6 and CBL may reflect feedback that restrains the initial signaling response. Thus, the later attenuation of the precursor program is compatible with an initiating period of excessive signaling. The thermal responses of several PKC isoforms and PRKD2 further implicate a broader DAG-responsive network, without assigning the phenotype to a single PKC isoform.

Interestingly, the previously reported transcriptional signature of primary MEF2D-r BCP-ALL provides further evidence of altered DAG-responsive signaling. *DGKI* was among the most strongly upregulated genes, accompanied by increased *DGKH, DGKD* and *RASGRP1* and reduced expression of the RAS/ERK negative regulators *RASA2*, *DUSP5* and *DUSP6* ^2^. Consistent with this pattern, our previously published transcriptomic resource also shows marked enrichment of *DGKI* RNA expression in MEF2D-r BCP-ALL cell lines, and our earlier proteomic analysis identified increased DGKH abundance in this subgroup ^12^. These findings suggest that regulation of DAG signaling is selectively remodeled in this subtype. As DGKs metabolize DAG to phosphatidic acid and can constrain the magnitude and duration of antigen-receptor signaling, altered DAG turnover could contribute to maintaining MEF2D-r cells within a permissive signaling range, whereas Bryo-1-mediated C1-domain activation may bypass or overwhelm this regulatory state and drive signaling beyond a tolerated threshold. The contribution of individual DGK isoforms to this putative signaling rheostat remains to be determined.

The NFATC1 phosphosite response provides an additional potential link between acute Ca²⁺-responsive signaling and remodeling of the precursor B-cell state. NFAT proteins are regulated by the balance between calcineurin-dependent dephosphorylation and opposing kinase activity, which controls their subcellular localization and transcriptional output ^18–20^. NFATC1 is also functionally implicated in early B-cell development, including regulation of EBF1 and formation of the pre-BCR-associated developmental state ^20^. The Bryo-1-induced changes in NFATC1 phosphorylation observed here are therefore compatible with engagement of a developmental Ca²⁺/NFAT axis alongside MEK-ERK signaling. However, NFATC1 localization, transcriptional activity and functional requirement were not directly tested, and these data do not establish NFATC1 as the determinant separating Bryo-1-sensitive from resistant cells.

The distribution of sensitivity argues for combining genetic classification with functional assessment. MEF2D rearrangement can enrich for responsive disease, but the responses outside this group suggest that the relevant signaling configuration can arise in other genetic backgrounds. Consistent with this possibility, PMA was recently reported to induce PKC-ERK activation, cell-cycle arrest, apoptosis and differentiation-associated changes in the Ph+ BCP-ALL cell line SUP-B15, suggesting that susceptibility to excessive DAG-responsive signaling is not restricted to MEF2D-r leukemia ^22^. Conversely, ERK activation in YCUB-4R was accompanied by limited cytotoxicity and recovery of CD179a and Ki-67. This pattern suggests that resistance may involve the ability to recover from an initial perturbation or to tolerate its downstream consequences. It also identifies a practical distinction between pathway engagement and commitment to cell death. A dynamic assessment incorporating early signaling and later persistence of the precursor program may therefore be more informative than an isolated pERK measurement. Together, these findings support a precision medicine framework in which MEF2D rearrangement enriches for Bryo-1 sensitivity, while functional response profiling may refine patient selection within and beyond the genetically defined subgroup.

The high-resolution phosphoproteomic analysis was performed in KASUMI-9 using 100 nM Bryo-1, whereas the phenotypic response was observed at substantially lower concentrations across multiple models. Although low dose ERK activation and MEK dependence support conservation of the core signaling response, whether the broader phosphosignaling architecture and its kinetics distinguish sensitive from resistant models remains to be determined.

The phenotypic changes further suggest that leukemia cells retain a capacity to alter their developmental state despite oncogenic transformation. Loss of surrogate light-chain components and IL7R, together with CD20 acquisition and reduced levels of several precursor-associated markers, is compatible with a maturation-associated shift. The heterogeneous trajectories of individual markers favor partial remodeling over a uniform transition to a mature B-cell phenotype. This distinction has therapeutic relevance: changing selected features of the leukemic state may be sufficient to alter treatment susceptibility, even without terminal differentiation. The response therefore provides a rationale for exploring how pharmacological manipulation of developmental programs can change the targets and dependencies available to subsequent therapies.

One potential translational application of this cell-state remodeling is combination with antigen-directed immunotherapy. Antigen density influences CAR activation and killing, with the recognition threshold depending on receptor design ^23^. Ramakrishna and colleagues showed that Bryo-1 increased CD22 levels and improved the activity and persistence of CD22-directed CAR-T cells in preclinical leukemia models ^15^. Subsequent work further demonstrated context-dependent modulation of CD19 and CD22 levels by Bryo-1 in pre-B ALL and showed that these effects can interact with CAR-T-mediated antigen modulation ^24^. The increases in CD19 and CD22 observed here, together with substantial CD20 induction in several models therefore do not by themselves establish a new immunotherapy mechanism, but show that immunotherapy-relevant antigen remodeling accompanies the broader developmental response in MEF2D-r leukemia. This provides a rationale to test whether the Bryo-1-induced MEF2D-r state alters susceptibility to CD19-, CD20- or CD22-directed therapies. Quantitative assessment of antigen abundance and its persistence after drug removal would help identify when Bryo-1-treated cells are most susceptible to each targeted modality.

The preservation of antigen induction in YCUB-4R is particularly relevant to this strategy. It demonstrates that a weak cytotoxic response does not necessarily eliminate the opportunity to modify the leukemia cell surface. We therefore propose that direct killing and antigen modulation should be evaluated as distinct therapeutic outputs. Whether this separable surface-remodeling phenotype translates into increased immune-mediated killing, particularly in MEF2D-r models with limited direct cytotoxicity, will require functional testing.

CD20 induction offers an additional route to clinical translation. CD20-directed treatment already has a clinical precedent in CD20-positive adult B-lineage ALL, including the randomized evaluation of rituximab with chemotherapy ^25^. Recent work reported that Bryo-1 enhanced CD20 CAR-T activity in preclinical lymphoma models in association with increased antigen levels and mitigation of trogocytosis-associated antigen loss ^13^. Thus, Bryo-1-mediated CD20 induction itself is not unique to MEF2D-r leukemia. Our findings instead place CD20 acquisition within the broader Bryo-1-induced remodeling of the MEF2D-r precursor state. The central question is whether the induced antigen is sufficiently abundant and durable to confer functional sensitivity to the chosen immune effector.

The immunotherapeutic consequences of Bryo-1 may also extend beyond target antigen abundance. Wang and colleagues reported increased susceptibility of pre-B-cell acute lymphoblastic leukemia (pre-B-ALL) cells to natural killer (NK)-like and antigen-independent killing by activated CAR-T cells after Bryo-1 treatment, with different effects in a Burkitt lymphoma model ^24^. Our current experiments do not directly address these functional immune effects, but the coordinated remodeling observed in MEF2D-r cells provides a rationale for testing whether antigen modulation translates into enhanced immune-mediated killing. Such studies should distinguish leukemia cell priming from direct effects of Bryo-1 on the immune effector, for example by comparing tumor-cell pretreatment followed by drug removal with concurrent exposure. More broadly, these observations raise the possibility that Bryo-1 could provide complementary therapeutic effects through direct leukemia-cell killing in responsive models, increased target-antigen expression that may render less cytotoxic responders more susceptible to antigen-directed cellular therapies, and modulation of immune-effector function.

The reduction in CXCR4 raises a further possibility involving the bone marrow microenvironment. CXCR4-CXCL12 interactions contribute to stromal protection of BCP-ALL cells, and experimental CXCR4 inhibition can enhance their sensitivity to chemotherapy ^26^. The rapid loss of surface CXCR4 and reduction in total CXCR4 abundance observed here therefore suggest that Bryo-1 could alter access to protective niche signals. Together with reduced IL7R, this provides a rationale to investigate whether the remodeled leukemia state becomes less supported by its environment. Stromal cocultures and *in vivo* trafficking studies could determine whether these changes translate into reduced protection or altered tissue distribution. Such an effect would add a microenvironmental component to the direct signaling response and antigen remodeling.

The concentration dependence of the response has important implications for dosing. Several models showed stronger inhibition at intermediate concentrations than at the highest concentrations tested, and this pattern was also evident in the SJBALL020987 and SJBALL020667 PDX-derived cells. Bryo-1 has previously shown isoform- and concentration-dependent effects on PKC regulation, including a biphasic effect on PKCδ downregulation in a different cellular system ^27^. Differential regulation of DAG-responsive proteins or downstream feedback could contribute to the response observed here, although the responsible mechanism remains unresolved. These findings favor selection of a biologically active exposure range rather than assuming that increasing the dose will increase efficacy. In KASUMI-9, a 5-minute exposure to 100 nM Bryo-1 was followed by a sustained reduction in live-cell numbers after washout. This observation supports investigation of intermittent schedules, while the exposure needed to reproduce this effect at lower concentrations remains to be defined.

Bryo-1 has a useful translational advantage because it has already been evaluated intravenously in multiple phase I and II clinical studies, including patients with lymphoid malignancies. Early clinical studies defined dosing schedules, demonstrated pharmacodynamic modulation of PKC and identified cumulative myalgia as a principal dose-limiting toxicity ^28,29^. Although clinical activity was limited in several unselected phase II oncology studies, this prior human experience provides a foundation for biomarker-guided re-evaluation in a newly defined responsive context. As an investigational agent, Bryo-1 would require a dedicated clinical protocol for BCP-ALL. The relevant opportunity is to use existing clinical knowledge to evaluate a newly defined responsive context, with schedules guided by leukemia pharmacodynamics and the intended combination. Low-nanomolar activity in culture is encouraging, but effective marrow exposure and tolerability must be linked experimentally. The observed antagonism with MEK inhibition further argues for combination partners that preserve the signaling response required for direct killing.

The *ex vivo* response in PDX-derived cells supports pursuing this translation beyond established cell lines. The next decisive studies should establish in vivo antileukemic activity and a therapeutic window relative to normal hematopoietic cells, alongside functional testing of the proposed immunotherapy combinations. For patient selection, short *ex vivo* assays could assess both loss of viability and induction of the intended immune target. These measurements address different treatment objectives and could inform whether Bryo-1 is best developed for direct cytoreduction, immune priming, or both. The substantial changes in immunophenotype also warrant consideration when designing residual-disease monitoring, because treated leukemia cells may no longer closely match their pretreatment marker profile.

These findings identify MEF2D-r BCP-ALL as a leukemia subtype with a therapeutically exploitable developmental signaling vulnerability. Activation of the PKC-associated signaling network engages an ERK-dependent program that compromises leukemia cell viability and disrupts the precursor state, revealing a potential treatment strategy for this high-risk subtype. More broadly, our results suggest that developmental signaling constraints retained by malignant precursor B cells can be therapeutically re-engaged to redirect leukemia cell fate. The accompanying remodeling of therapeutically relevant surface antigens, including in a poorly cytotoxic responder, represents an additional and potentially exploitable consequence of this reprogramming that warrants further evaluation in combination with immunotherapy.

## Methods

### Cell lines and cell culture

93 BCP-ALL cell lines were maintained either in RPMI 1640 medium containing 2 mM stable glutamine (L-Ala-L-Gln dipeptide; Sigma-Aldrich), supplemented with 10% fetal bovine serum (FBS; Sigma-Aldrich), 20 mM HEPES (Gibco), 1 mM sodium pyruvate (Sigma-Aldrich), 1× MEM non-essential amino acids (Sigma-Aldrich), and 1× penicillin-streptomycin (Sigma-Aldrich), or in Iscove’s Modified Dulbecco’s Medium (IMDM; Sigma-Aldrich) supplemented with 10% or 20% FBS and 1× penicillin-streptomycin. Cells were cultured at 37°C in a humidified atmosphere with 5% CO₂. Cell line origins and corresponding culture media are provided in **Table S1**. All cell lines were routinely tested for mycoplasma contamination using the MycoAlert Mycoplasma Detection Kit (Lonza) and authenticated by short tandem repeat (STR) profiling (Eurofins Genomics).

### PDX samples

PDX samples were obtained from St. Jude Children’s Research Hospital Public Resource of Patient-derived and Expanded Leukemias (PROPEL; https://propel.stjude.cloud). Basic information for the PDX models is provided in **Table S2**. PDX expansion and drug screening were performed as described earlier ^30,31^. In brief, PDX sample were thawed and cells were intravenously injected into 6-8 week old immunodeficient male or female NSG mice (Jackson Laboratory, #005557). Leukemia cells were isolated from the spleen or bone marrow of the mice and were used for a short-term functional *ex vivo* assay ^30,31^. All animal experiments were performed in compliance with the regulatory guidelines under protocol number: 2024-32-Grundantrag and were approved by the animal research institute (ZETT) at Heinrich Heine University Düsseldorf.

### Drug sensitivity assay

Cells were resuspended in culture medium and seeded at 1 × 10⁶ cells/mL in 96-well flat-bottom white plates (Thermo Fisher Scientific; Cat. No. 136101). DMSO levels were normalized across the plate with a maximum final concentration of 0.1%. Bryo-1 was serially diluted to final concentrations ranging from 0.0015 to 100 nM in a 96-well plate format. Cells were incubated with Bryo-1 for 3 days or 6 days, after which cell viability was measured using CellTiter-Glo 2.0 ATP reagent (Promega; G9242) on a Victor 3V plate reader or ApoTox-Glo Triplex assay (Promega; G6321).

### Reagents and antibodies

Bryostatin-1 (203811) was purchased from Merck (Sigma Aldrich). Trametinib (16292) and U0126 (70970) were obtained from Cayman Chemical. Primary antibodies for Western blot included ERK1/2 (4695; Cell Signaling Technology), Phospho-ERK1/2 (Thr202/Tyr204) (4370; Cell Signaling Technology), MEF2D (25651; Cell Signaling Technology), PRKCD (PA5-81985; Invitrogen), β-actin (sc-47778; Santa Cruz Biotechnology), Phospho-ZAP70 (ab194800; Abcam), ZAP70 (2709S; Cell Signaling Technology), SYK (80460S; Cell Signaling Technology) and SOD1 (sc-17767; Santa Cruz Biotechnology). Secondary antibodies used for Western blot detection included IRDye 680RD goat anti-mouse IgG (926-68070), IRDye 680RD goat anti-rabbit IgG (926-68071), IRDye 800CW goat anti-mouse IgG (926-32210), and IRDye 800CW goat anti-rabbit IgG (926-32211), all purchased from LI-COR Biosciences.

### Flow Cytometry analysis

Cells were collected at the indicated time points, washed with phosphate-buffered saline (PBS), and stained with Zombie NIR Fixable Viability Dye (BioLegend) for dead-cell exclusion, followed by staining with fluorochrome-conjugated antibodies against the indicated surface markers. Cells were subsequently fixed and permeabilized using the eBioscience Foxp3/Transcription Factor Staining Buffer Set (Thermo Fisher Scientific) according to the manufacturer’s instructions and stained intracellularly for CD179a and Ki-67. Samples were acquired on a Sony ID7000 Spectral Cell Analyzer (Sony Biotechnology). Single-stained controls were used to generate spectral reference spectra, and spectral unmixing was performed using the ID7000 system software.

Unmixed FCS files were imported into FlowJo v10.10.2 (BD Life Sciences). Acquisition-related anomalies were identified and removed using the flowAI plugin before downstream analysis. Cells were sequentially gated based on forward and side scatter, singlets, and Zombie NIR-negative viable cells. Marker expression was quantified as indicated in the corresponding figures.

### Western blot and isothermal dose-response Cellular Thermal Shift Assay (ITDR-CETSA)-Western blot

KASUMI-9, YCUB-4 and YCUB-4R cells were treated with 100nM Bryo-1 for time points (0h, 2h, 24h, 48h). Cells were collected, denatured in SDS sample buffer, separated on NuPAGE 4-12% Bis-Tris gels, and transferred to PVDF membranes. Membranes were blocked in 5% non-fat dry milk in TBST and incubated overnight at 4 °C with antibodies against ERK1/2, p-ERK1/2, MEF2D, or SOD-1, followed by HRP-conjugated secondary antibody. Bands were detected using Clarity ECL substrate on an iBright CL1000 Imaging System.

KASUMI-9 cells were treated with 1.5 nM Bryo-1 for 0, 15, and 30 min, and 2, 8, and 24 h. Cells were collected, denatured in SDS sample buffer, separated on NuPAGE 4-12% Bis-Tris gels, and transferred to PVDF membranes. Membranes were blocked using LI-COR blocking buffer (LI-COR Biosciences) and incubated overnight at 4 °C with primary antibodies against phospho-ERK1/2 (Thr202/Tyr204) and total ERK1/2. Membranes were subsequently incubated with IRDye 680RD- and IRDye 800CW-conjugated secondary antibodies (LI-COR Biosciences), and fluorescence signals were acquired using an Odyssey infrared imaging system (LI-COR Biosciences). GAPDH was detected using a directly rhodamine-conjugated anti-GAPDH recombinant Fab fragment (clone AbD22549, catalog no. 12004167; Bio-Rad), without secondary antibody incubation, and imaged using an iBright FL1500 Imaging System (Thermo Fisher Scientific). Band intensities were background-corrected, and phospho-ERK1/2 and total ERK1/2 signals were individually normalized to GAPDH before calculating the phospho-ERK1/2-to-total ERK1/2 ratio.

Cells were washed twice with HBSS and lysed in 100 mM HEPES and 20 mM MgCl₂ by five freeze-thaw cycles. Lysates were aliquoted and treated for 10 min with Bryo-1 at 0, 4, 10, 20, 30, 50, 100, or 500 nM in DMSO. Samples were heated at 48 °C for 3 min and centrifuged at 14,000 × g for 20 min at 4 °C. Supernatants were collected, denatured in SDS sample buffer, separated on NuPAGE 4-12% Bis-Tris gels, and transferred to nitrocellulose membranes. Membranes were blocked in 5% non-fat dry milk in TBST and incubated overnight at 4 °C with antibodies against PRKCD, followed by HRP-conjugated secondary antibody (SCBT, sc-2005). Bands were detected using Clarity ECL substrate on an iBright CL1000 Imaging System.

### 2DTPP

KASUMI-9 cells were subjected to two-dimensional thermal proteome profiling (2DTPP) ^16,17,32^ using Bryo-1 treatment in intact cells. Cells were treated with 0, 4, 20, 100, or 500 nM bryostatin-1 for 2 h at 37 °C, washed twice with HBSS, and resuspended in HBSS. For each drug concentration, samples were aliquoted and heated for 3 min at 42, 45, 48, 51, 54, 57, 60, 63, or 66 °C. In-cell samples were subsequently lysed by five freeze-thaw cycles. Insoluble material was removed by centrifugation at 14,000 × g for 20 min, and soluble protein fractions were collected. Proteins were reduced with TCEP, alkylated with chloroacetamide, and sequentially digested with LysC and trypsin. Peptides were labeled with TMTpro 16plex reagents, with the nine temperatures divided across three TMT sets (42-48, 51-57, and 60-66 °C). Samples within each set were pooled, desalted, and further fractionated by high-pH reversed-phase chromatography, with concatenation into 19 fractions per TMT set.

Fractions were analyzed using an UltiMate 3000 RSLCnano system coupled to a Q Exactive HF mass spectrometer (Thermo Fisher Scientific). Peptides were separated on a 50-cm C18 column using a 110-min gradient at 0.25 µL/min. MS1 spectra were acquired at 60,000 resolution over m/z 300-1500, followed by data-dependent HCD MS/MS of the five most intense precursors at 30,000 resolution and a normalized collision energy of 30%. Precursors were isolated using a 1.2 m/z window, with AGC targets of 1 × 10⁶ and 1 × 10⁵ and maximum injection times of 100 ms for MS1 and MS2, respectively.

Raw files were processed using the IsobarQuant pipeline and searched with Mascot v2.6 against the Swiss-Prot human proteome database. Carbamidomethylation of cysteine and TMTpro labeling of lysine were specified as fixed modifications, whereas N-terminal acetylation, methionine oxidation, and N-terminal TMTpro labeling were variable modifications. Searches were performed using trypsin/P specificity, up to three missed cleavages, precursor and fragment mass tolerances of 10 ppm and 0.05 Da, respectively, and peptide-spectrum matches were used for TMTpro-based protein quantification.

### Quantitative proteomics

Quantitative proteomics was performed as previously described ^33^. KASUMI-9 cells were treated with 100 nM Bryo-1 or vehicle (DMSO) for 6 or 24 h. Cells were washed twice with HBSS and lysed in 4% SDS, 25 mM HEPES (pH 7.6), and 1 mM DTT. Lysates were heated at 95 °C for 5 min, sonicated, and clarified by centrifugation at 14,000 × g for 15 min. Protein concentrations were determined using the DC protein assay, and 190 µg protein per sample was alkylated with chloroacetamide and processed using the single-pot solid-phase-enhanced sample preparation (SP3) method. Proteins were sequentially digested with LysC and trypsin at enzyme-to-protein ratios of 1:50 overnight at 37 °C. Peptides were quantified, dried, resuspended in 0.1 M triethylammonium bicarbonate, and labeled with TMTpro reagents for 2.5 h. Labeled samples were pooled at equal ratios and desalted using Strata-X-C cartridges.

For deep proteome fractionation, pooled peptides were separated by high-resolution isoelectric focusing (HiRIEF)^34^ using 24-cm immobilized pH-gradient strips (pH 3-10). Peptides were passively eluted into 72 fractions, dried, and analyzed by LC-MS/MS using an UltiMate 3000 RSLCnano system coupled to a Q Exactive HF mass spectrometer (Thermo Fisher Scientific). Peptides were separated on a 50-cm C18 column using a 110-min gradient and analyzed in data-dependent acquisition mode. MS1 spectra were acquired at 60,000 resolution over m/z 300-1500, followed by HCD MS/MS of the five most intense precursors at 30,000 resolution with a normalized collision energy of 30%.

Raw files were processed using the ddamsproteomics v3.0 workflow. MS data were converted using ProteoWizard v3.0.24172 and searched with MS-GF+ v2023.01.1202 against the Ensembl release 113 Homo sapiens protein database. Searches used a precursor mass tolerance of 10 ppm, fully tryptic peptides, carbamidomethylation of cysteine as a fixed modification, oxidation of methionine as a variable modification, and TMTpro modifications on lysine residues and peptide N-termini. Peptide-spectrum matches were rescored using Percolator v3.5, and identifications were controlled at 1% false discovery rate (FDR). TMTpro reporter-ion quantification was performed using OpenMS v3.1.0, with HiRIEF fraction information incorporated during identification. Protein-level quantification was based on reporter-ion signals from peptides assigned uniquely to genes and normalized across samples. Hardklör v2.3.2 was used for feature detection, and downstream protein-level statistical analysis was performed using DEqMS v1.18.0.

### Phosphoproteomics

KASUMI-9 cells were used to generate four replicates and treated with 100 nM Bryo-1 for 0, 15, 45, or 120 min at 37 °C. Cells were washed twice with HBSS and lysed in 4% SDS, 25 mM HEPES (pH 7.6), and 1 mM DTT. Lysates were heated at 95 °C for 10 min, sonicated, clarified by centrifugation at 20,000 × g for 20 min, and protein concentrations were determined using the DC protein assay. For each sample, 400 µg protein was alkylated with chloroacetamide and processed using the SP3 method. Proteins were sequentially digested with LysC and trypsin overnight at 37 °C and peptides were labeled with TMTpro reagents for 2 h. Labeled samples were pooled at equal ratios, desalted using Strata-X-C cartridges, and fractionated by high-pH reversed-phase chromatography into 96 fractions, which were concatenated to 26 fractions. Five percent of each fraction was retained for total-proteome analysis, while the remaining material was combined into 13 fractions for phosphopeptide enrichment.

Phosphopeptides were enriched using Ti²⁺-IMAC magnetic beads. Peptides were dissolved in 80% acetonitrile, 6% TFA, and 0.1 M lactic acid and incubated with equilibrated beads for 45 min at 30 °C. After washing, phosphopeptides were eluted with 5% NH₄OH into 10% TFA, dried, and analyzed by LC-MS/MS using an UltiMate 3000 RSLCnano system coupled to a Q Exactive HF mass spectrometer (Thermo Fisher Scientific). Peptides were separated on a 50-cm C18 column using a 150-min gradient at 0.25 µL/min. MS1 spectra were acquired at 60,000 resolution over m/z 300-1500, followed by data-dependent HCD MS/MS of the five most intense precursors at 30,000 resolution and normalized collision energy of 30%. Precursors were isolated using a 2 m/z window, with AGC targets of 1 × 10⁶ and 2 × 10⁵ and maximum injection times of 100 and 400 ms for MS1 and MS2, respectively.

The non-enriched total-proteome fractions were analyzed using the ddamsproteomics v3.0 workflow. Raw files were converted using ProteoWizard v3.0.24172 and searched with MS-GF+ v2023.01.1202 against the Ensembl release 113 Homo sapiens protein database. Searches used fully tryptic peptides, a precursor mass tolerance of 10 ppm, carbamidomethylation of cysteine as a fixed modification, oxidation of methionine as a variable modification, and TMTpro modifications on lysine residues and peptide N-termini. Peptide-spectrum matches were rescored using Percolator v3.5, reporter-ion quantification was performed using OpenMS v3.1.0, and protein-level statistical analysis used DEqMS v1.18.0.

Enriched phosphopeptide data were reanalyzed using FragPipe v23.1 with MSFragger v4.3 against a reviewed human UniProt proteome database supplemented with contaminants and decoy sequences. Searches used strict tryptic specificity with up to two missed cleavages, precursor and fragment mass tolerances of 20 and 10 ppm, respectively, and peptide lengths of 7-50 amino acids. Carbamidomethylation of cysteine and TMTpro labeling of lysine residues and peptide N-termini were specified as fixed modifications, whereas methionine oxidation and phosphorylation of serine, threonine, and tyrosine were variable modifications. PSMs were rescored using Percolator v3.7.1, phosphorylation sites were localized using PTMProphet v7.3.0, and identifications were filtered at a 1% FDR. Protein inference and FDR control were performed using Philosopher v5.1.2. TMTpro reporter-ion intensities were extracted using IonQuant v1.11.11 and summarized using TMT-Integrator v6.1.1; phosphosite-level quantification required a minimum site-localization probability of 0.75.

### Databases and bioinformatic analyses

Differential protein abundance was calculated using DEqMS ^35,36^. Differentially abundant proteins in KASUMI-9 cells were defined by |log₂ fold change (FC) | > 0.5 and adjusted *p*-value < 1 × 10⁻⁴. Over-representation pathway enrichment analysis was performed using enrichR^37^. Phosphoproteomic comparisons were performed using limma ^38^. Kinase activity inference was performed using KSEA ^39^. Site-specific PTM signature enrichment analysis (PTM-SEA) of phosphoproteomics data sets was performed using ssGSEA2 ^40^. Figures were created using BioRender, GraphPad Prism and Adobe Illustrator.

## Data Availability

The mass spectrometry proteomics data will be deposited to the ProteomeXchange Consortium via the PRIDE partner repository.

## Code Availability

The R analysis code will be available on GitHub.

## Acknowledgements

This study was supported by grants from the Swedish Childhood Cancer Foundation (R.J., TJ2016-0035, PR2016-0019, PR2019-0025, PR2022-0009 and PR2024-0065); The Swedish Cancer Society (R.J., 23-2843 Pj and 24-0866 SIA); the Dr. Åke Olsson Foundation for Hematological Research (R.J., 2021-00130); Cancer Research Foundations of Radiumhemmet (R.J., 221132 and 241243); and KI Research Foundation Grants (X.Q., 2024-02353); R.J. acknowledges the Karolinska Institutet and Science for Life Laboratory. The authors would like to thank the Biomedicum Flow Cytometry Core Facility (BFC), and in particular Alda Saldan, for their support with flow cytometry experiments and data acquisition. The authors would like to thank Rubin Joshi and Takahiro Seki for their assistance during the project. The authors would like to thank Monique L. den Boer and Judith Boer from Princess Máxima Center for Pediatric Oncology, Utrecht, The Netherlands for providing the M4A1-M2B9 cell line. Figures 3A, 4C and 4F were created in BioRender. Proteomics, L. (2026) https://BioRender.com/harksw7.

## Author Contributions

R.J. designed and coordinated the study together with Q.G. and X.Q.. R.J. and J.L. provided funding, resources and access to the mass spectrometry platform. Q.G., X.Q. I.R.L. and F.P. developed the methodology and performed mass spectrometry experiments with help from G.M. and O.H., Q.G. performed and analyzed drug screening together with X.Q., N.S. and T.E.. K.A. and T.I. provided cell lines and K.P.T. provided clinical samples. M.K. and S.B. performed PDX expansion and ex vivo drug sensitivity testing together with Q.G.. Q.G., X.Q. and L.A. developed data analysis strategies and analyzed the data with help from I.R.L., F.P.. Q.G. and X.Q. drafted the manuscript with input from R.J. All authors read and approved the current version of the manuscript.

## Competing Interests

The authors declare no competing financial or non-financial interests.

## Notes

### Competing Interest Statement

The authors have declared no competing interest.

