## Supplementary_figures for "Bryostatin-1 exploits a developmental signaling vulnerability in MEF2D-rearranged B-cell precursor acute lymphoblastic leukemia"

**Supplementary Figures 1-4 and Table 1-2 related to:**

**Bryostatin-1 exploits a developmental signaling vulnerability in MEF2D-rearranged B-cell precursor acute lymphoblastic leukemia**

**Authors**

Quentin Glaziou^1§^, Xuekang Qi^1§^, Luay Aswad^1^, Isabelle Rose Leo^1^, Frederik Post^1^, Marie Kemkes^2^, Melina Vogt^2^, Owen F.J. Hovey^1^, Georgios Mermelekas^1^, Nona Struyf^3^, Tom Erkers^3^, Katja Pokrovskaja Tamm^4^, Koshi Akahane^5^, Takeshi Inukai^5^, Sanil Bhatia^2^, Janne Lehtiö^1^, Rozbeh Jafari^1*^

**Affiliations**

^1^ Clinical Proteomics Mass Spectrometry, Department of Oncology-Pathology, Karolinska Institutet, Science for Life Laboratory, Tomtebodavägen 23A, 171 65 Solna, Sweden

^2^ Department of Pediatric Oncology, Hematology and Clinical Immunology, Medical Faculty, Heinrich Heine University Düsseldorf, Germany.

^3^ Molecular Precision Medicine, Department of Oncology-Pathology, Karolinska Institutet, Science for Life Laboratory, Tomtebodavägen 23A, 171 65 Solna, Sweden

^4^ Department of Oncology-Pathology, Karolinska Institutet, J6:140 BioClinicum, Akademiska stråket 1, 171 64 Solna, Sweden

^5^ Global Leukemia Cell-line Assembly Network, Department of Pediatrics, School of Medicine, University of Yamanashi, Chuo, Japan

^§^ These authors contributed equally.

^*^ Corresponding author

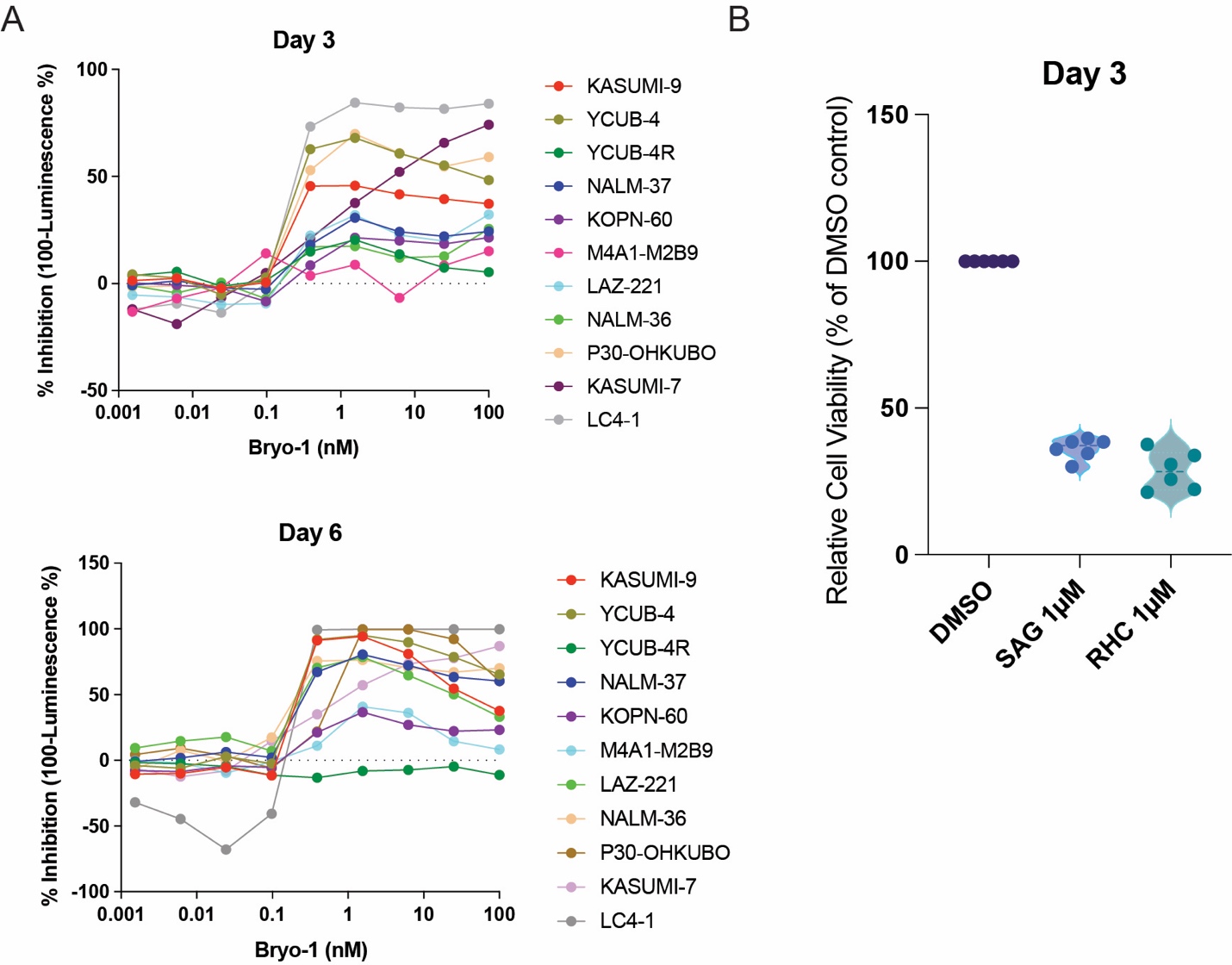

**Supplementary Figure 1. Concentration dependence and DAG-associated features of the Bryo-1 response.**

(**A**) Concentration-response profiles in indicated cell lines following 3 or 6 days of treatment with the indicated concentrations of Bryo-1. ATP-based viability was measured using CellTiter-Glo, and inhibition was calculated as 100 - (100 × luminescence in treated cells/luminescence in corresponding DMSO controls). Negative inhibition values indicate luminescence above the DMSO control. Colors identify individual cell lines. (**B**) Relative viability of KASUMI-9 cells after 3 days of treatment with 1 µM 1-stearoyl-2-arachidonoyl-sn-glycerol (SAG) or 1 µM RHC 80267 (RHC), a diacylglycerol lipase inhibitor. Viability was measured using CellTiter-Glo and expressed as a percentage of the corresponding DMSO-treated controls. Violin plots show the distribution of measurements, with individual points representing six independent biological experiments.

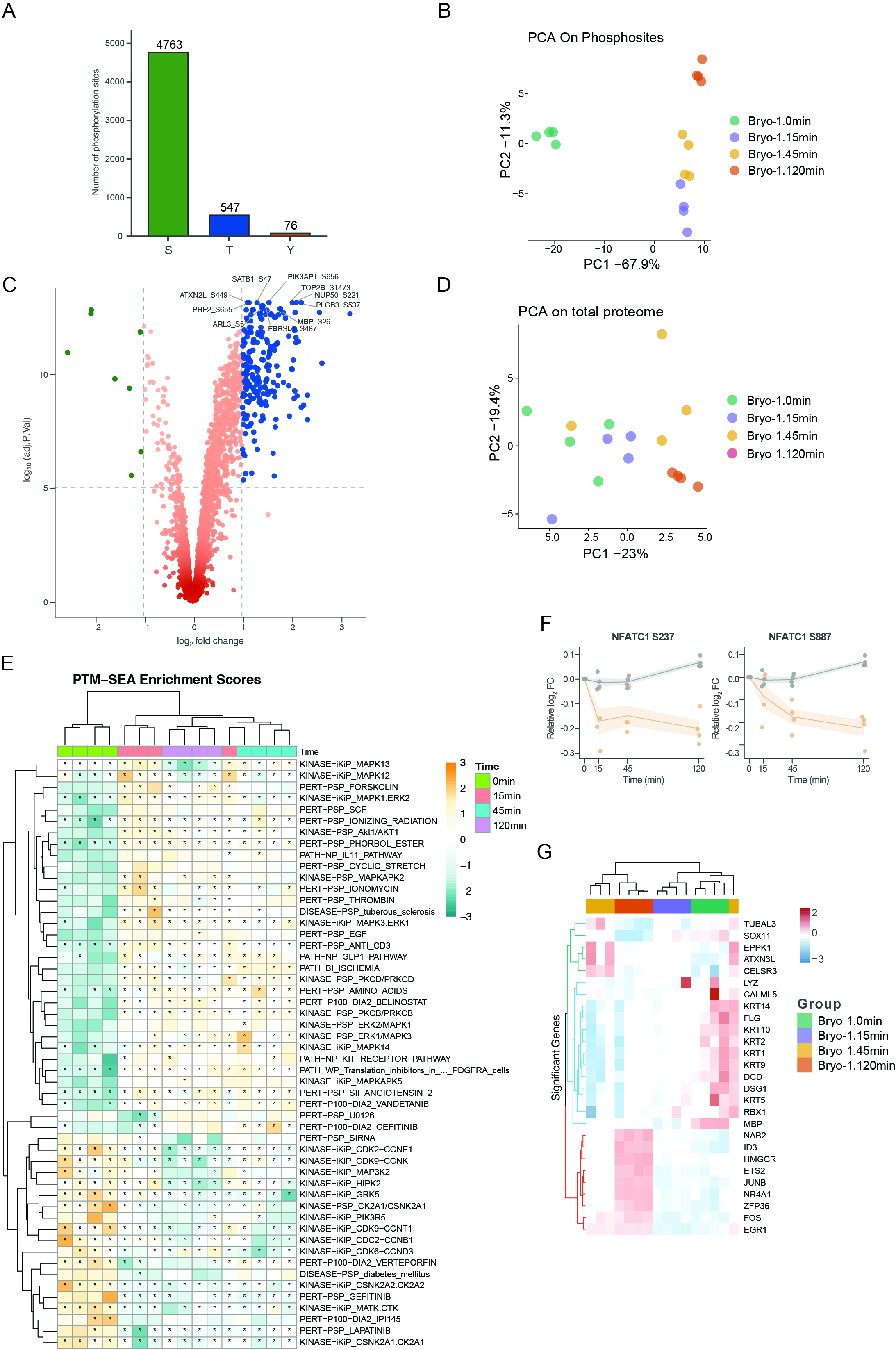

**Supplementary Figure 2. Global phosphoproteomic remodeling and NFATC1 phosphoregulation following Bryostatin-1 treatment.**

**(A)** Distribution of quantified phosphorylation sites according to phosphorylated residue in the KASUMI-9 time-course phosphoproteomic dataset (n=4 biological replicates per time point). A total of 4,763 serine, 547 threonine and 76 tyrosine class I phosphorylation sites were quantified. **(B)** Principal component analysis (PCA) of phosphosite abundances in KASUMI-9 cells at baseline and after 15, 45 or 120 min of treatment with 100 nM Bryo-1. Each point represents an individual biological replicate and colors indicate treatment time. PC1 and PC2 account for 67.9% and 11.3% of the total variance, respectively. **(C)** Volcano plot of phosphosite changes following 15 min Bryo-1 treatment relative to baseline. Each point represents an individual phosphosite; the x-axis indicates log2 fold change and the y-axis indicates -log10 adjusted *p*-value. Selected strongly regulated phosphosites are annotated. Dashed lines indicate the effect-size and statistical thresholds used for highlighting regulated sites. **(D)** PCA of the matched total-proteome measurements obtained from the phosphoproteomic experiment. Each point represents an individual biological replicate and colors indicate treatment time. PC1 and PC2 account for 23.0% and 19.4% of the total variance, respectively. **(E)** Post-translational modification signature enrichment analysis (PTM-SEA) across the 0-120 min Bryo-1 time course. Rows represent phosphosite signatures and columns represent individual biological replicates grouped by treatment time. Enrichment scores are standardized within each row. Asterisks denote signatures meeting the indicated false-discovery threshold. **(F)** Temporal regulation of NFATC1 phosphosites S237 and S887 following treatment with 100 nM Bryo-1. Phosphosite abundance is shown as relative log2 fold change from baseline together with the corresponding total NFATC1 protein abundance. Points represent biological replicates and lines summarize the time-dependent trajectories. **(G)** Hierarchical clustering of proteins showing temporal abundance changes across the matched 0-120 min total-proteome time course. Rows represent proteins and columns represent individual samples, with abundance values standardized within each row. Colors in the upper annotation indicate treatment time.

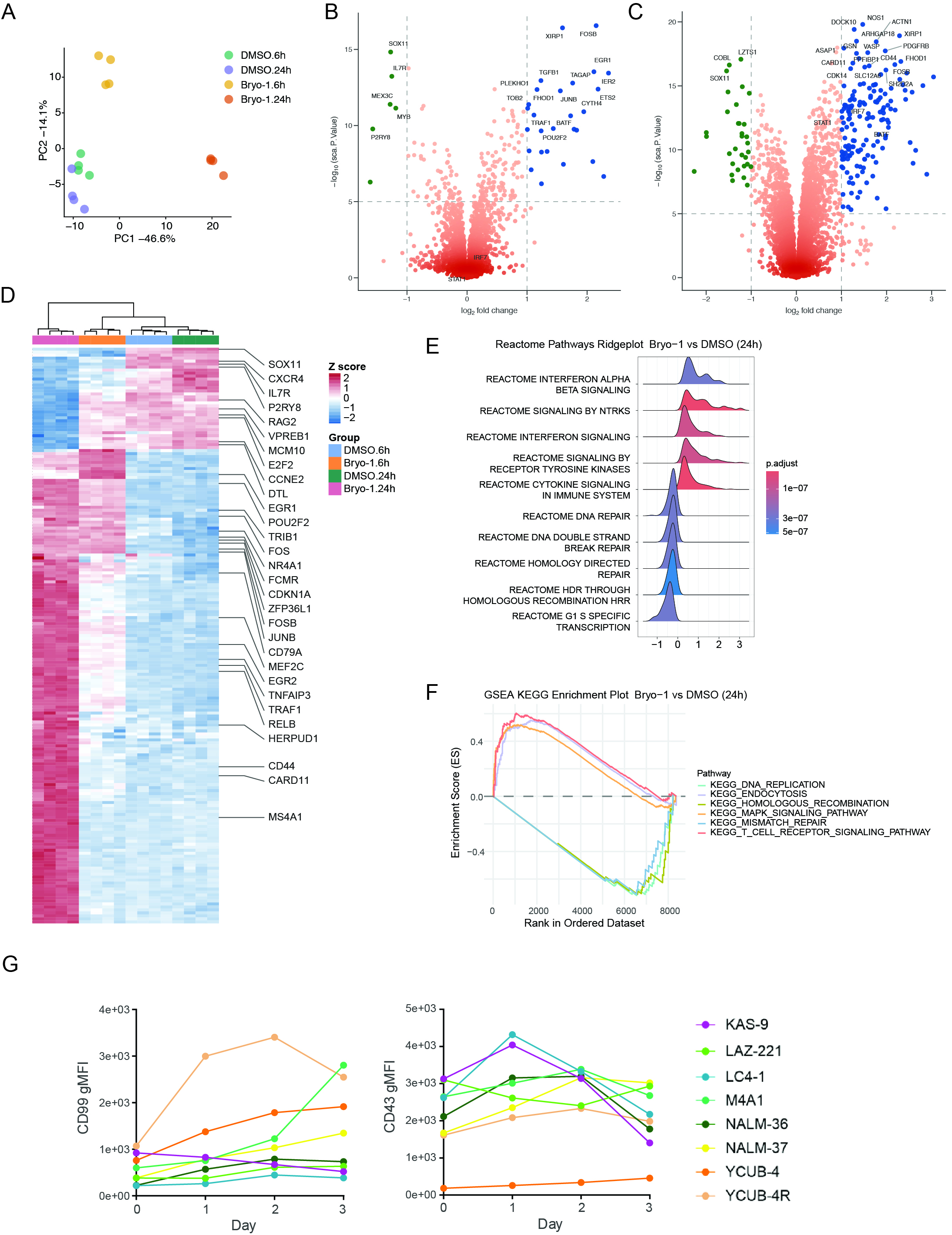

**Supplementary Figure 3. Temporal proteome remodeling following Bryostatin-1 treatment.**

**(A)** Principal component analysis (PCA) of the KASUMI-9 proteome following 6 or 24 h of treatment with 100 nM Bryo-1 or corresponding time-matched DMSO controls. Each point represents an individual biological replicate and colors indicate treatment condition, n=4 biological replicates per time point. PC1 and PC2 account for 46.6% and 14.1% of the total variance, respectively. **(B, C)** Differential protein abundance following Bryo-1 treatment for **(B)** 6 h and **(C)** 24 h relative to the corresponding time-matched DMSO controls. Each point represents one quantified protein; the x-axis indicates log2 fold change and the y-axis indicates -log10 DEqMS *p*-value. Proteins meeting the indicated statistical and effect-size thresholds are highlighted, and selected regulated proteins are annotated. **(D)** Hierarchical clustering of proteins showing significant abundance changes across the 6- and 24-h proteome. Rows represent proteins and columns represent biological replicates from DMSO and Bryo-1-treated conditions. Protein abundances are standardized within each row (z-score). Selected proteins associated with the precursor B-cell state, proliferation, immediate-early signaling and immune regulation are annotated. **(E)** Reactome gene-set enrichment analysis of the 24-h Bryo-1 versus DMSO proteome. Ridge plots show the distribution of ranked protein-level changes for selected enriched pathways; color indicates the adjusted enrichment *p*-value. Positive enrichment includes interferon- and cytokine-associated signaling programs, whereas negatively enriched categories include DNA repair, homologous recombination and G1/S-associated transcription. **(F)** Gene-set enrichment analysis of selected KEGG pathways at 24 h after Bryo-1 treatment relative to time-matched DMSO controls. Running enrichment scores are plotted across the ranked proteome. Selected pathways illustrate coordinated changes in DNA replication and repair, MAPK signaling and immune/signaling programs. **(G)** Time courses of surface CD99 and CD43 expression in the indicated BCP-ALL cell lines following treatment with 100 nM Bryo-1. Expression was measured by flow cytometry and is presented as geometric mean fluorescence intensity (gMFI), with day 0 corresponding to the pretreatment baseline. Colors identify individual cell lines (n=1).

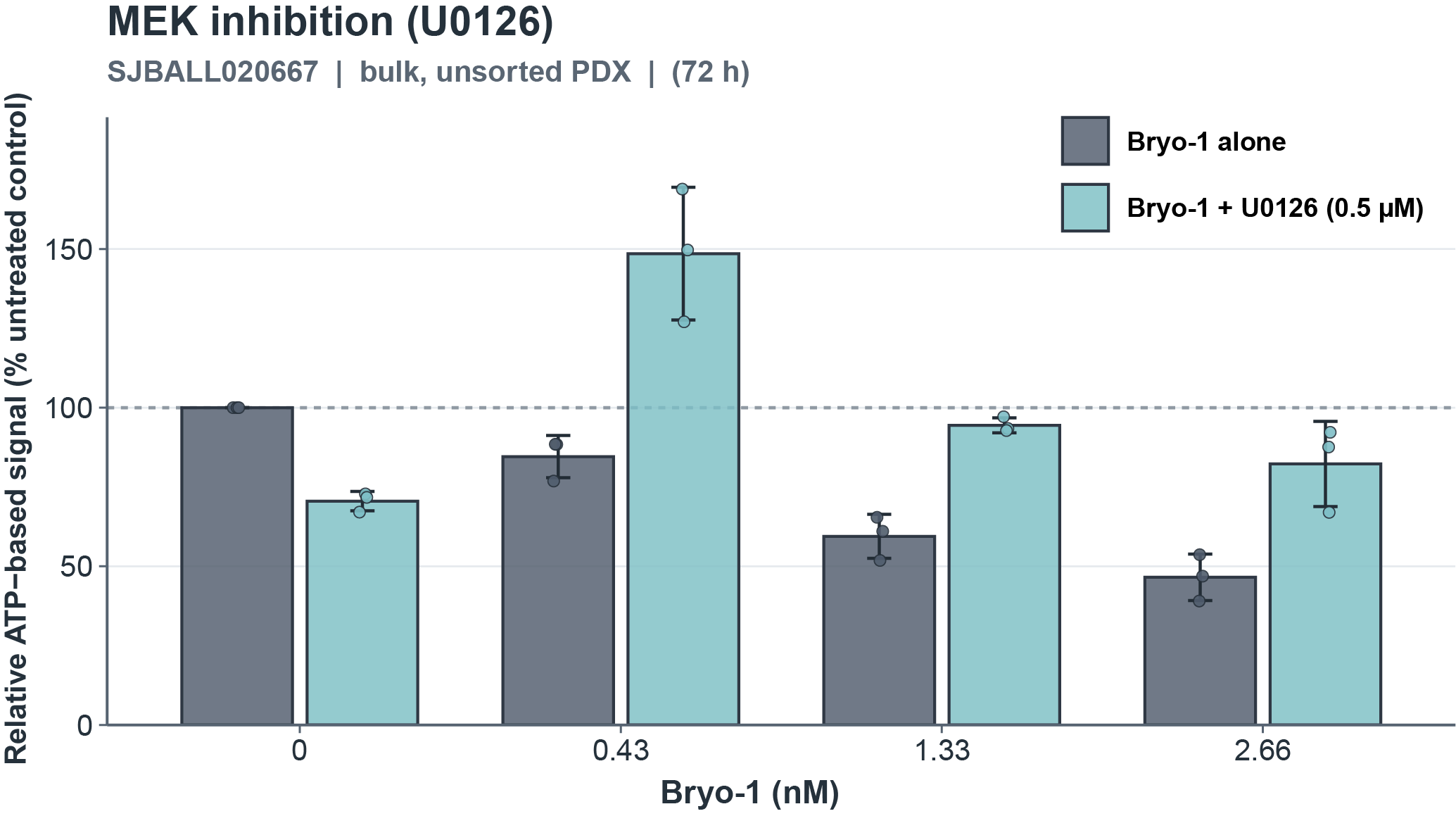

**Supplementary Figure 4. MEK inhibition attenuates the Bryo-1 associated loss of viable cell signal in PDX-derived leukemia cells.** ATP-based viability measurements in bulk, unsorted SJBALL020667 (*MEF2D::BCL9*) PDX-derived cells following 72 h of treatment with indicated concentrations of Bryo-1 alone or in combination with 0.5 µM U0126 (MEK inhibitor). CellTiter-Glo luminescence was normalized to the untreated control (0 nM Bryo-1, 0 µM U0126), set to 100%. Bars represent the mean ± SD of three technical wells per condition, with individual dots indicating each measurement. The dashed horizontal line indicates the untreated control level (100%).

**Supplementary Table 1. Cell lines used in this study**

| **Cell Line Name** | **Subtype** | **Grouped_Subtype** | **Tissue** | **Age** | **Sex** | **Time** | **Relapse** | **Medium**  **(medium name+ FBS%)** | **Cellosaurus_ID** |
| --- | --- | --- | --- | --- | --- | --- | --- | --- | --- |
| **380** | *IGH::MYC* | MYCr | PB | 15 | M | PD | 1st | RPMI.10 | CVCL_1047 |
| **697** | *TCF3::PBX1* | TCF3r | BM | 12 | M | PD | 1st | RPMI.10 | CVCL_0079 |
| **ALL-PO** | *KMT2A::AFF1* | KMT2Ar | BM | 0.33 | F | Dx | Diagnosis | RPMI.10 | CVCL_1069 |
| **COG-LL-319h** | *TCF3::PBX1* | TCF3r | BM | 13 | F | Dx | Diagnosis | IMDM.20 | CVCL_M161 |
| **COG-LL-355h** | *ETV6::RUNX1* | ETV6r | BM | 7 | M | Dx | Diagnosis | IMDM.20 | CVCL_M164 |
| **COG-LL-394h** | *ABL1::ZMIZ1* | ABL1r | BM | NA | F | PD | NA | IMDM.20 | NA |
| **COG-LL-402h** | *TCF3::PBX1* | TCF3r | BM | 9 | M | Dx | NA | IMDM.20 | NA |
| **HAL-01** | *TCF3::HLF* | TCF3r | PB | 17 | F | PD | Relapse | RPMI.10 | CVCL_1242 |
| **KASUMI-10** | *KMT2A::MLLT1* | KMT2Ar | PB | 0.5 | F | Dx | NA | IMDM.20 | CVCL_8485 |
| **KASUMI-2** | *TCF3::PBX1* | TCF3r | BM | 15 | M | Dx | Relapse | RPMI.10 | CVCL_0590 |
| **KASUMI-9** | *MEF2D::HNRNPUL1* | MEF2Dr | PB | 19 | M | NA | Relapse | RPMI.10 | CVCL_8488 |
| **KOPN-8** | *KMT2A::MLLT1* | KMT2Ar | PB | 0.25 | F | Dx | Diagnosis | RPMI.10 | CVCL_1866 |
| **LC4-1** | *MEF2D::HNRNPUL1* | MEF2Dr | PB | 13 | F | Dx | Diagnosis | RPMI.10 | CVCL_1374 |
| **MHH-CALL-2** | *B::Other* | B-Other | PB | 15 | F | Dx | Diagnosis | RPMI.20 | CVCL_1409 |
| **MHH-CALL-3** | *TCF3::PBX1* | TCF3r | BM | 11 | F | Dx | Diagnosis | RPMI.20 | CVCL_0089 |
| **MHH-CALL-4** | *IGH::CRLF2* | CRLF2 | PB | 10 | M | Dx | Diagnosis | RPMI.10 | CVCL_1410 |
| **NALL-1** | *PAX5::ETV6* | PAX5r | PB | 14 | M | PD | 1st | RPMI.10 | CVCL_3039 |
| **NALM-16** | *B::Other* | B-Other | PB | 12 | F | PD | 1st | RPMI.10 | CVCL_1834 |
| **NALM-6** | *DUX4::IGH* | DUX4-IGH | PB | 19 | M | PD | 1st | RPMI.10 | CVCL_0092 |
| **P30-OHKUBO** | *MEF2D::HNRNPUL1* | MEF2Dr | BM | 11 | F | PD | 3rd | RPMI.10 | CVCL_1631 |
| **RCH-ACV** | *TCF3::PBX1* | TCF3r | BM | 8 | F | Dx | Relapse | RPMI.10 | CVCL_1851 |
| **REH** | *ETV6::RUNX1* | ETV6r | PB | 15 | F | PD | 1st | RPMI.10 | CVCL_1650 |
| **SEM** | *KMT2A::AFF1* | KMT2Ar | PB | 5 | F | PD | 1st | RPMI.10 | CVCL_0095 |
| **SUP-B15** | *BCR::ABL1* | ABL1r | BM | 9 | M | PD | 2nd | RPMI.10 | CVCL_0103 |
| **TANOUE** | *IGH::MYC* | MYCr | PB | 11 | M | NA | NA | RPMI.10 | CVCL_1852 |
| **TMD5** | *BCR::ABL1* | ABL1r | PB | 19 | M | Dx | Diagnosis | AMEM.10 | CVCL_3218 |
| **BALM-14** | *IGH::MYC* | MYCr | BM | 13 | M | NA | NA | IMDM.20 | CVCL_5692 |
| **BALM-19** | *TCF3::HLF* | TCF3r | BM | 25 | M | Dx | NA | IMDM.20 | CVCL_5695 |
| **BALM-26** | *TCF3::HLF* | TCF3r | BM | 25 | M | PD | NA | IMDM.20 | CVCL_5702 |
| **BALM-9** | *IGH::MYC* | MYCr | PB | 12 | F | NA | NA | RPMI.10 | CVCL_5707 |
| **COG-LL-528h** | *IGH::MYC* | MYCr | BM | NA | NA | Dx | NA | IMDM.20 | NA |
| **KASUMI-7** | *MEF2D::HNRNPUL1* | MEF2Dr | PB | 29 | F | NA | Relapse | RPMI.10 | CVCL_8486 |
| **KASUMI-8** | *BCR::ABL1* | ABL1r | PB | 48 | M | NA | Relapse | IMDM.20 | CVCL_8487 |
| **KASUMIA-541** | *NUP214::ABL1* | ABL1r | PB | 26 | M | NA | NA | RPMI.20 | CVCL_8523 |
| **LAZ-221** | *MEF2D::BCL9* | MEF2Dr | PB | 24 | F | Dx | Diagnosis | IMDM.20 | CVCL_1830 |
| **MUTZ-5** | *B::Other* | B-Other | PB | 26 | M | PD | NA | IMDM.20 | CVCL_1873 |
| **NAGL-1** | *CRLF2::IGH* | CRLF2 | BM | 18 | F | NA | NA | RPMI.10 | CVCL_3038 |
| **NALM-19** | *KMT2A::AFF1* | KMT2Ar | PB | 26 | M | Dx | Diagnosis | IMDM.20 | CVCL_1835 |
| **NALM-20** | *BCR::ABL1* | ABL1r | PB | 62 | M | Dx | Diagnosis | RPMI.20 | CVCL_1836 |
| **NALM-21** | *BCR::ABL1* | ABL1r | BM | 63 | M | PD | 1st | RPMI.10 | CVCL_1837 |
| **NALM-33** | *IGH::MYC* | MYCr | PB | 72 | M | PD | NA | RPMI.20 | CVCL_5671 |
| **PALL-2** | *BCR::ABL1* | ABL1r | PB | 45 | M | NA | NA | IMDM.20 | CVCL_3122 |
| **RS4-11** | *KMT2A::AFF1* | KMT2Ar | BM | 32 | F | PD | 1st | RPMI.10 | CVCL_0093 |
| **M4A1-M2B9** | *MEF2D::BCL9* | MEF2Dr | BM | 15 | M | Dx | Diagnosis | IMDM.20 | NA |
| **BALM-18** | *IGH::MYC* | MYCr | PB | 35 | M | NA | NA | IMDM.20 | CVCL_5694 |
| **BALM-22** | *TCF3::HLF* | TCF3r | BM | 25 | M | NA | NA | IMDM.20 | CVCL_5699 |
| **BALM-25** | *B::Other* | B-Other | BM | 59 | M | NA | NA | IMDM.20 | CVCL_5701 |
| **TOM-1** | *BCR::ABL1* | ABL1r | BM | 54 | F | NA | NA | IMDM.20 | CVCL_1895 |
| **NALM-29** | *BCR::ABL1* | ABL1r | PB | 46 | M | NA | NA | IMDM.20 | CVCL_5667 |
| **NALM-36** | *MEF2D::BCL9* | MEF2Dr | PB | 37 | F | NA | NA | IMDM.20 | CVCL_5674 |
| **KOPN61** | *MEF2D::BCL9* | MEF2Dr | NA | 6 | NA | Dx | Diagnosis | IMDM.10 | CVCL_A331 |
| **KOPN71** | *MEF2D::BCL9* | MEF2Dr | NA | 11 | NA | Dx | Diagnosis | IMDM.10 | NA |
| **KOPN39** | *MEF2D::BCL9* | MEF2Dr | NA | 9 | NA | Dx | Diagnosis | IMDM.20 | CVCL_W273 |
| **KOS20** | *MEF2D::BCL9* | MEF2Dr | NA | 14 | NA | PD | Relapse | IMDM.20 | NA |
| **KOPN70** | *MEF2D::BCL9* | MEF2Dr | NA | 9 | NA | PD | Relapse | IMDM.20 | NA |
| **KOPN41** | *ETV6::RUNX1* | ETV6r | NA | 4 | NA | Dx | NA | IMDM.10 | CVCL_3943 |
| **KOPN79** | *ETV6::RUNX1* | ETV6r | NA | 3 | NA | PD | Relapse | IMDM.10 | CVCL_A334 |
| **KOPN46** | *MEF2D::HNRNPUL1* | MEF2Dr | NA | 9 | NA | Dx | Diagnosis | IMDM.10 | CVCL_W274 |
| **KOCL77** | *KMT2A::AFF1* | KMT2Ar | NA | 1 | NA | PD | Relapse | IMDM.20 | NA |
| **KOCL69** | *KMT2A::AFF1* | KMT2Ar | NA | 1 | NA | PD | Relapse | IMDM.20 | CVCL_3995 |
| **KOPN60** | *MEF2D::DAZAP1* | MEF2Dr |  | 14 | NA | Dx | Relapse | IMDM.20 | CVCL_A330 |
| **KOCL33** | *KMT2A::MLLT1* | KMT2Ar | NA | 1 | NA | Dx | Diagnosis | IMDM.20 | CVCL_3991 |
| **KOCL51** | *KMT2A::MLLT1* | KMT2Ar | NA | 1 | NA | Dx | Diagnosis | IMDM.20 | CVCL_6865 |
| **KOPN62** | *MEF2D::HNRNPUL1* | MEF2Dr | NA | 72 | NA | NA | NA | IMDM.20 | CVCL_A332 |
| **KOPN84** | *B::Other* | B-Other | NA | 15 | NA | PD | Relapse | IMDM.20 | CVCL_A335 |
| **KOPN49** | *B::Other* | B-Other | NA | 12 | NA | PD | Relapse | IMDM.20 | CVCL_W276 |
| **KOPN85** | *B::Other* | B-Other | NA | 6 | NA | PD | Relapse | IMDM.20 | NA |
| **KOPN32** | *B::Other* | B-Other | NA | 1 | NA | PD | Relapse | IMDM.20 | CVCL_A325 |
| **YAMN90R** | *TCF3::PBX1* | TCF3r | NA | 4 | NA | PD | Relapse | IMDM.20 | NA |
| **KOCL50** | *KMT2A::MLLT1* | KMT2Ar | NA | 1 | NA | PD | Relapse | IMDM.20 | CVCL_6866 |
| **KOPN66bi** | *BCR::ABL1* | ABL1r | NA | 10 | NA | PD | Relapse | IMDM.20 | CVCL_7979 |
| **YAMN96** | *MEF2D::HNRNPUL1* | MEF2Dr | NA | 8 | NA | Dx | Diagnosis | IMDM.20 | NA |
| **KOCL45** | *KMT2A::AFF1* | KMT2Ar | NA | 1 | NA | PD | Relapse | IMDM.10 | CVCL_3993 |
| **KOCL44** | *KMT2A::MLLT1* | KMT2Ar | NA | 1 | NA | PD | Relapse | IMDM.10 | CVCL_3992 |
| **PRE-ALP** | *TCF3::PBX1* | TCF3r | BM | 6 | NA | Dx | Diagnosis | IMDM.10 | NA |
| **NALM-27** | *BCR::ABL1* | ABL1r | NA | 38 | NA | Dx | Diagnosis | IMDM.20 | NA |
| **YACL95** | *KMT2A::MLLT3* | KMT2Ar | NA | 1 | NA | Dx | Diagnosis | IMDM.20 | NA |
| **YAMN92** | *TCF3::PBX1* | TCF3r | NA | 14 | NA | Dx | Relapse | IMDM.20 | CVCL_A080 |
| **KOPN72bi** | *BCR::ABL1* | ABL1r | NA | 9 | NA | PD | NA | IMDM.20 | CVCL_7981 |
| **KOPB26** | *KMT2A::MLLT3* | KMT2Ar | PB | 1 | NA | PD | Relapse | IMDM.10 | CVCL_7972 |
| **KOPN57bi** | *BCR::ABL1* | ABL1r | BM | 11 | NA | Dx | Diagnosis | IMDM.20 | CVCL_7978 |
| **KOPN35** | *KMT2A::MLLT10* | KMT2Ar | PB | 2 | NA | PD | Relapse | IMDM.20 | CVCL_A327 |
| **YCUB-2** | *TCF3::HLF* | TCF3r | PB | 4 | M | PD | Relapse | IMDM.20 | CVCL_A082 |
| **YCUB-4** | *MEF2D::BCL9* | MEF2Dr | PB | 7 | M | Dx | Diagnosis | IMDM.20 | CVCL_A083 |
| **YCUB-4R** | *MEF2D::BCL9* | MEF2Dr | PB | 7 | M | PD | Relapse | IMDM.20 | CVCL_A084 |
| **YCUB-5** | *P2RY8::CRLF2* | CRLF2 | BM | 12 | M | Dx | Diagnosis | IMDM.20 | CVCL_A085 |
| **YCUB-6** | *TCF3::PBX1* | TCF3r | PB | 12 | M | PD | Diagnosis | IMDM.20 | CVCL_A087 |
| **BEL-1** | *KMT2A::AFF1* | KMT2Ar | PB | 41 | F | PD | NA | IMDM.20 | CVCL_0081 |
| **BALM-23** | *TCF3::HLF* | TCF3r | BM | 24 | M | Dx | Diagnosis | IMDM.10 | CVCL_5700 |
| **NALM-24** | *BCR::ABL1* | ABL1r | PB | 42 | F | NA | NA | IMDM.10 | CVCL_3888 |
| **BALM-24** | *IGH::MYC* | MYCr | PB | 46 | F | NA | NA | IMDM.10 | CVCL_B3M5 |
| **NALM-37** | *MEF2D::BCL9* | MEF2Dr | BM | 37 | F | PD | 1st | IMDM.10 | CVCL_5676 |
| **NALM-30** | *BCR::ABL1* | ABL1r | BM | 39 | M | PD | 1st | IMDM.10 | CVCL_5668 |

Abbreviations: BM, bone marrow; PB, peripheral blood; Dx, diagnosis; PD, progressive disease; F, female; M, male; NA, not available; MEF2Dr, MEF2D-rearranged; KMT2Ar, KMT2A-rearranged; TCF3r, TCF3-rearranged; ABL1r, ABL1-rearranged; ETV6r, ETV6-rearranged; PAX5r, PAX5-rearranged; MYCr, MYC-rearranged.

**Supplementary Table 2. Patient-derived xenograft (PDX) models used in this study**

| **Sample ID** | **Leukemia subtype** | **Genomic Driver** | **Species Engrafted** | **Age at Diagnosis (Years)** | **Age Group** | **Disease status at sampling** | **Publications (PMID:)** |
| --- | --- | --- | --- | --- | --- | --- | --- |
| SJBALL020987_D1 | MEF2D | *MEF2D::BCL9* | NSG | 18 | AYA | Diagnosis | 27824051 |
| SJBALL020667_D1 | MEF2D | *MEF2D::BCL9* | NSG | 14 | Pediatric | Diagnosis | 27824051 |
| SJBALL021005_D1 | MEF2D | *MEF2D::BCL9* | NSG | 17 | AYA | Diagnosis | 27824051 |

Abbreviations: AYA, adolescent and young adult; NSG, NOD-scid IL2Rγnull; PDX, patient-derived xenograft.
